# Using a non-destructive sugar-feeding assay for sporozoite detection and estimating the extrinsic incubation period of *Plasmodium falciparum* in mosquito vectors

**DOI:** 10.1101/2020.12.03.408435

**Authors:** Edwige Guissou, Jessica L. Waite, Matthew Jones, Andrew S. Bell, Eunho Suh, Koudraogo B. Yameogo, Nicaise Djegbe, Dari F. Da, Domonbabele FdS Hien, Rakiswende S. Yerbanga, Anicet G. Ouedraogo, Kounbobr R. Dabiré, Anna Cohuet, Matthew B. Thomas, Thierry Lefèvre

## Abstract

Despite its epidemiological importance, the time *Plasmodium* parasites take to achieve development in the vector mosquito (the extrinsic incubation period, EIP) remains poorly characterized. A novel non-destructive assay designed to estimate EIP in single mosquitoes, and more broadly to study *Plasmodium* – *Anopheles* vectors interactions, is presented. The assay uses small pieces of cotton wool soaked in sugar solution to collect malaria sporozoites from individual mosquitoes during sugar feeding to monitor infection status over time. This technique has been tested across four natural malaria mosquito species of Africa and Asia, six parasite isolates of *Plasmodium falciparum*, and across a range of temperatures relevant to malaria transmission in field conditions. We find that monitoring individual infectious mosquitoes is feasible, although due to the frequency of mosquito sugar feeding and inter-individual variation in infection intensity, there is inherent risk that this technique will result in some false negatives. The sensitivity rate ranged from 0.27 to 0.81 depending on mosquito species and on infection intensity in mosquitoes used to collect saliva. Using this non-destructive technique, the estimated median extrinsic incubation period of *P. falciparum* at 27°C was 11 to 14 days depending on mosquito species and parasite isolate. Long-term individual tracking also revealed that sporozoite transfer onto cotton wool can occur at least until day 40 post-infection. In addition to contributing to a better understanding of EIP and mosquito to human transmission with implications for improving epidemiological models, this technique also allows to link different transmission traits at the mosquito individual level. As one example, we found a significant relationship between EIP and mosquito lifespan, with short individual EIP associated with short mosquito lifespan. Correlations between mosquito/parasite traits often reveal trade-offs and constraints and have important implications for understanding the evolution of parasite transmission strategies.

## Introduction

The extrinsic incubation period (EIP) of *Plasmodium* is the duration of the parasite’s development within the mosquito that starts with the ingestion of gametocytes (the human to mosquito transmission stage) in a blood meal and ends with the sporozoite invasion of the salivary glands when the mosquito becomes infectious (Lefevre et al., 2018; Ohm et al., 2018). Following the consumption of gametocyte-positive blood, the male and female gametocytes first egress from the enveloping erythrocytes and fertilize in the mosquito midgut. Within a few hours, the resulting zygotes complete a meiosis and develop into motile ookinetes that cross the midgut wall and lodge to the basal lamina of the midgut where they mature into oocysts. Within the oocysts, mitotic divisions produce large number of sporozoites, which break out into the mosquito body cavity, and invade the mosquito salivary glands at least one week post-infection (depending on the malaria species (Collins et al., 1969; Knowles and Basu, 1943; Nikolaev, 1935) and temperature (Detinova, 1963; Ohm et al., 2018; Paaijmans et al., 2012, 2010; Shapiro et al., 2017). EIP plays a crucial role in malaria vectorial capacity (a standard measure of malaria transmission potential), as small changes in EIP can have a large effect on the number of mosquitoes living long enough to be able to transmit parasites (Brady et al., 2016; MacDonald, 1957; Smith and McKenzie, 2004). The earlier the parasite invades mosquito salivary glands, the sooner the vector becomes infectious, and will have the potential to infect more humans over its lifespan.

Despite such epidemiological importance, the estimation of EIP has been limited to a small number of vector-parasite combinations, and sources of variability in EIP remain largely unknown. Besides temperature and parasite species, evidence that EIP can be influenced by other genetic and environmental factors, and possibly be associated with other key transmission parameters such as mosquito longevity, is limited (Lefevre et al., 2018; Ohm et al., 2018). The reason for this gap of knowledge probably lies in the difficulty of measuring EIP in the laboratory. Ohm et al. (Ohm et al., 2018) showed that EIP50 (the time until 50 percent of infected mosquitoes become infectious, which equates to the median time of sporozoite development) is the most useful measure for transmission modeling, but finding the midpoint of a transmission curve is labor intensive. Current methods consist in experimentally infecting a large batch of mosquitoes and dissecting a number of salivary glands at several time points (e.g. every day or two) to monitor parasite maturation. This process suffers several limitations. First, it requires infecting several hundreds of mosquitoes at the same time. This number increases if there is not an already well-defined time range for the EIP of the studied parasite species, or when infection prevalence is low. Second, it requires skilled researchers to dissect mosquito salivary glands and examine them for sporozoites under high magnification (400-1000x). Third, this approach results in an estimation of EIP at the mosquito population level, hence preventing the characterization of direct associations between EIP and other life-history traits such as mosquito longevity, fecundity, or parasite density. Faced with these limitations, other researchers have tried to solve the problems by developing new methods. These methods consist in marking parasites with a green fluorescent protein (GFP) to allow parasites to be seen without necessarily dissecting the mosquitoes. However, this also limits the scope of the work of using experimentally infected mosquitoes with these GFP strains only, and is still technically complex (Ramakrishnan et al., 2012; Vaughan et al., 2012). As an alternative to mosquito dissection, Billingsley et al. (Billingsley et al., 1991) developed an antibody based blot assay to see if sporozoites were deposited with mosquito saliva on a sugar substrate, their successes suggesting this technique could be further developed.

In the context of large-scale vector-borne disease surveillance and elimination programs, some systems have been developed whereby the sugar feeding behavior of mosquito vectors is similarly exploited to detect pathogens. As during blood feeding, mosquito vectors salivate during sugar feeding, not only to facilitate ingestion by diluting concentrated nectars, but also to initiate sugar digestion (Stone and Foster, 2013). While salivating, pathogens can be expelled. New field surveillance systems rely on traps where mosquitoes are provided access to sugar-baited substrates from which expelled pathogens can then be detected using qRT-PCR. Such sugar-based surveillance systems to track pathogens in wild mosquito populations have now been successfully applied to arboviruses (Burkett-Cadena et al., 2016; Flies et al., 2015; Girod et al., 2016; Hall-Mendelin et al., 2010; Johnson et al., 2015; Lothrop et al., 2012; Ritchie et al., 2013; van den Hurk et al., 2014) and are currently being developed for malaria (Brugman et al., 2018; Melanson et al., 2017).

Pathogen detection through mosquito sugar feeding has also recently been used in laboratory settings for investigating mosquito-virus interactions. Applications include the potential use of *Wolbachia* to reduce dengue transmission (Ye et al., 2015), the effect of *Aedes aegypti* genetic variation on the EIP of a dengue virus (Ye et al., 2016), the temporal dynamic of mosquito competence for dengue and Chikungunya viruses (Alto et al., 2017; Fontaine et al., 2016), the changes in viral population structure over time (Grubaugh et al., 2017), mosquito immune response to Chikungunya and Zika viruses (Zhao et al., 2018) and characterizing the EIP of a West Nile Virus in *Culex tarsalis* (Danforth et al., 2018). Repeated sampling using this non-destructive method requires fewer mosquitoes and offers a unique opportunity to investigate the dynamic nature of vector-pathogen interactions.

In this study, a novel non-destructive assay is described based on the exploitation of mosquito sugar feeding to reveal infection with malaria parasites and designed to study *Plasmodium* – *Anopheles* vectors interactions. In particular, we provide proof-of-concept that this assay can be used to estimate *Plasmodium* EIP in single mosquitoes across a range of different vector species, parasite strains and densities, and temperatures.

## Methods

### Mosquito colonies and maintenance

Experiments were conducted between two labs, the one at the Institut de Recherche en Sciences de la Santé (IRSS, Bobo Dioulasso, Burkina Faso), and the other at Pennsylvania State University (PSU, Pennsylvania, USA), and used five mosquito colonies in total.

1. *Anopheles gambiae* (IRSS), obtained from an outbred colony brought into the lab in 2008 and repeatedly replenished with F1 from wild-caught mosquito females collected in Soumousso, (11°23’14“N, 4°24’42“W), 40 km from Bobo Dioulasso, and identified by routine SINE PCR (Santolamazza et al., 2008).
2. *Anopheles coluzzii* (IRSS), obtained from an outbred colony brought into the lab in 2016 from wild-caught mosquito females collected in Kou Valley (11°24’N, 4°24’59’’W), 30 km from Bobo Dioulasso, and identified by SINE PCR (Santolamazza et al., 2008).
3. *Anopheles arabiensis* (IRSS) obtained from an outbred colony brought into the lab in 2015 from wild-caught gravid females collected in Soumousso, and identified by routine SINE PCR (Santolamazza et al., 2008).
4. *Anopheles stephensi* (PSU) established in the lab from eggs shipped from Walter Reed (USA) in June 2015.
5. *An. gambiae/An. coluzzi* hybrid form (PSU) established in the lab from eggs shipped from NIH from the G3 line in August 2015 (G3 strain hereafter).

All mosquitoes were maintained under standard conditions (12/12 hrs daylight/darkness light cycle, 27 ± 2°C, RH 70% ± 10%). IRSS colonies were maintained by blood meals either on anesthetized rabbit hosts (protocol was approved by both the Office of Laboratory Animal Welfare of US Public Health Service (Assurance Number: A5928-01) and national committee of Burkina Faso (IRB registration #00004738 and FWA 00007038), animals were cared for by trained personnel and veterinarians) or at PSU by using bell jar feeders with either sausage casing or parafilm to hold a human blood meal mixed with CPDA-1 as an anticoagulant (sourced from either Valley Biomedical, Winchester, VA, USA or Biological Specialty Corp. Colmar, PA, USA). 10% (w/v) glucose sugar solution was provided for colony sugar feeding, with 0.05% of para aminobenzoic acid (PABA) at PSU and without PABA at IRSS.

### Infectious feeds

At IRSS, mosquito females were infected by using blood from naturally gametocyte-infected patients recruited in villages surrounding Bobo Dioulasso, Burkina Faso, using direct membrane feeding assays as previously described (Hien et al., 2016; Ouédraogo et al., 2013). Ethical approval was obtained from the institutional ethical review committee under agreement no 2017-003/MESRSI/CNRST/IRSS/CEIRES. At PSU mosquitoes were fed on blood infected with NF54 *P. falciparum* parasite cultures, (provided by the Johns Hopkins Malaria Institute Core Facility, NF54 obtained from MRA-1000, MR4, ATCC® Manassas Virginia), or were fed with NF54 *P. falciparum* wild type parasites cultured in the PSU lab following protocols described in (Shapiro et al., 2017) and provided by the Center for Infectious Disease Research, Seattle, WA. Blood for culturing and infectious feeds was sourced from either Valley Biomedical (Winchester, VA) or Biological Specialty Company (Colmar, PA). Gametocyte cultures reached approximately 2-4% mature gametocytemia and were between 14-17 days post gametocyte induction when cultures were diluted with a mixture of freshly washed red blood cells and inactivated human serum and fed to 3-5 day old mosquitoes. Mosquitoes were maintained as in (Shapiro et al., 2017) at 27 °C and after the 20 minute infectious feed were immediately moved to incubators set to experimental temperatures at 80% rH with +/− 0.5C accuracy and +/−5% rH. Data loggers were used in each incubator and checked to confirm the accuracy of the incubators and adjust any offsets as required prior to the start of the experiments; monitoring continued for the duration of the experiments. Mosquitoes were provided with a sugar meal of 10% glucose on cotton wool following feeding.

### Mosquito dissections

Mosquitoes were dissected and midguts examined under 100x to 400x total magnification under phase contrast and/or dark field for microscopic examination of oocysts in dissected midguts (either stained with 1% mercurochrome at IRSS, or unstained at PSU) from 7-9 days post-blood meal (dpbm). Sporozoites were observed in dissected salivary glands at 1000x total magnification, also under phase contrast from days 12-14 at 27°C unless other temperatures were specified.

### Salivary Pathogens In Transmission (“SPIT”) assay development

The assay used in the following experiments was optimized prior to use. Details about the assay development and optimization is provided in **Supplementary data 1**. This optimization process includes choosing the optimal substrate, minimum number of mosquito tested, and assay duration (S1, 1), DNA extraction method (S1, 2), and testing qPCR detection methods (S1, 3). Additionally, a standard curve was generated using final methods, and aspects of sample degradation were tested (S1, 4). All following experiments used the following optimized assay conditions: 15mg cotton wools soaked in 10% glucose solution (either with or without PABA added) were left overnight on 28 ml plastic drosophila tubes hosting single mosquitoes. The cottons were collected the following morning, and DNA extracted using a slightly modified protocol provided in the Qiagen DNeasy Blood and Tissue kit. After DNA precipitation using absolute ethanol, the cotton wool substrate was transferred and only removed after DNA elution. Testing with known quantities of sporozoites under various conditions showed the malaria DNA can degrade with exposure to heat and humidity, so it is best practice to collect the sporozoite-containing cotton wool samples daily and store frozen until ready for DNA extraction (S1, 4). DNA amplification and visualization using qPCR SYBR methods that produced the best results are provided next; all other supporting information is provided in **Supplementary data 1**. Additionally, the utility of the assay at different temperatures was tested to determine if transmission to the sugar saturated cotton might occur at 12 °C, 14 °C, 16 °C, 18 °C, 32 °C, and 34 °C using sporozoite stage infected mosquitoes. At least some positive samples were detected at all temperatures except 12 °C, suggesting that if mosquitoes are active and sugar feeding, this assay could be functional; details provided in **Supplementary data 2**.

### *qPCR detection of* Plasmodium falciparum

Detection and quantification of the sporozoites in cottons and in mosquito females used to collect saliva on cottons was performed by real-time PCR using standard procedures. We targeted the mitochondrial gene that codes for the cytochrome C Oxidase (Cox1) (Boissière et al., 2013). The sequences of the primers used were: qPCR-PfF 5’-TTACATCAGGAATGTTATTGC-3 ’and qPCR-PfR 5’-ATATTGGATCTCCTGCAAAT-3’. Sample reaction occurred in a total volume of 10 μl containing 1 μl of DNA (~ 40 ng/μl); 4.6μL of water; 2μl of 1x HOT Pol Eva Green qPCR Mix Plus ROX and 1.2μl of each primer at 5μM. The amplification began with an activation step at a temperature of 95 °C for 15 min and 40 cycles of denaturation at 95 °C to 15 s, annealing / extension at 58 °C for 30 s. Samples (cottons and female’s head/thorax) were considered positive for *P. falciparum* when the qPCR yielded a Cp < 35 and a 75 > Tm < 80 (at IRSS) or Cp < 35 and a 73 > Tm < 75 (at PSU).

### Experiment 1: Comparing estimates of the parasite’s EIP using the classic dissection approach versus the non-destructive individual “spit” assay

At IRSS, *An. gambiae* mosquitoes were exposed to an infectious blood meal from two *P. falciparum* isolates (A=168 gametocytes/ul of blood and B=1 208 gametocytes/ul of blood) using direct membrane feeding assay. Mosquitoes were kept at 27 °C and 75 % relative humidity, and the extrinsic incubation period of *P. falciparum* was assessed through both microscopic observation of mosquito gut and salivary glands from 8 to 16 dpbm (i.e. the classic destructive approach) and the individual “spit” assay (non-destructive approach) over the same period of time.

#### 1.1 Destructive approach: mosquito dissection and microscopic observation

From 8 to 16 dpbm, 8 to 20 mosquito females (median 14) exposed to isolate A or B were dissected daily. The presence and number of oocysts in mosquito guts and the sporozoites in the salivary glands were assessed microspically. Oocyst rupture in mosquito midgut and sporozoite invasion of salivary glands is highly asynchronous. For instance at 12 dpbm, while some oocysts are intact and keep developing within a given mosquito gut, others have already ruptured and released their sporozoites (**Supplementary data 3**). To assess the timing of sporozoite dissemination in mosquito salivary, three metrics were derived from the microscopic observation:

i. the proportion of infected mosquitoes with ruptured oocysts at 8-16 dpbm. This is the number of mosquitoes with at least one ruptured oocyst in their midguts out of the total number of infected mosquitoes (i.e. harboring either intact and/or ruptured oocysts);
ii. the proportion of ruptured oocysts at 8-16 dpbm. This is, for each infected mosquito, the number of ruptured oocysts out of the total number of oocysts (intact + ruptured);
iii. the proportion of oocyst-infected mosquitoes with sporozoites in their salivary glands at 8-16 dpbm. This is the number of oocyst-infected mosquitoes harboring sporozoites in their head/thoraces out of the total number of infected mosquitoes (i.e. harboring either intact and/or ruptured oocysts). Following microscopic observation, salivary glands were stored at −20°C in individual 1.5 ml Eppendorf tube for further qPCR determination of infection status; such that we were able to compare microscopic and molecular diagnostic for this metric.

#### 1.2 Non-destructive approach: the individual “Spit” assay

Twenty one (21) females per parasite isolate (A or B) were individually placed in 28 ml plastic drosophila tubes at 7 dpbm and cotton balls (15 mg/piece) soaked with 10% glucose solution were placed on the tube gauze at 17:00 until collection at 7:00 in sterile 1.5 ml Eppendorf tubes stored at −20 °C for further qPCR analysis. Cottons were replaced daily from 7 to 15 dpbm and saliva was collected from 8 to 16 dpbm. At the end of the experiment, the presence of parasites in the heads/thoraxes of the females used to collect mosquito saliva was tested using qPCR as described above. Individual EIP was defined as the time between the infectious blood meal and the first day of positive molecular detection by qPCR of *P. falciparum* from a cotton wool substrate collected for a given female.

### Experiment 2: Individual estimation of EIP in different mosquito species

#### 2A IRSS

Mosquito females of *An. gambiae*, *An. coluzzii*, and *An. arabiensis* were all exposed to an infectious blood meal from one volunteer carrying gametocytes (isolate C: 3024 gametocytes / ul of blood) using direct membrane feeding assays. At 7 dpbm, 20 females from each species were individually placed in plastic drosophila tubes for saliva collection using the spit assay as described above. In this experiment, saliva was collected daily from 8 to 14 dpbm and every other day from 14 to 20 dpbm and stored at −20°C before DNA was processed.

#### 2B PSU

From 6 dpbm cotton wool substrate was provided the same way as in experiment 2A to 15 individuals of *An. stephensi*. Cotton balls were left on cups from 14:30 until collection at 09:30 and replaced daily from 7-21 dpbm. Cotton balls were collected and stored for up to 1 week at −20 °C before transferring to −80 °C until DNA isolation and qPCR.

### Experiment 3: Determination of mosquito sugar feeding rate

#### 3A IRSS

##### Assay 1

Four-day-old females of *An. gambiae*, *An. coluzzi* and *An. arabiensis* received an uninfectious meal from rabbit blood. Two days later, 30 females from each species were placed in 180 ml plastic cups with white filter paper at the bottom and kept at 27 °C and 75% relative humidity. Every evening at 17:00 cotton balls soaked with 10% glucose and containing a blue dye (1 mg mL−1 Fast Green FCF/Xylene cyanole FF; Sigma-Aldrich) (Ignell et al., 2010) were deposited. The day after at 07:45, the cottons were removed from the plastic cups. At 16:00, the presence or absence of blue droppings on the paper was observed. The presence of dejection on the paper indicates that a sugar intake by the mosquito occurred and the absence of dejection is considered us no sugar intake. The same day at 17:00, new cottons were added to the same plastic cups, but with a 10% (w/v) glucose solution containing a yellow dye (1 mg mL−1 Acid Yellow 17; Sigma-Aldrich, St Louis, Missouri). This procedure was repeated during 8 consecutive days with the papers changed every day and blue and yellow dyes on cottons switched between days.

##### Assay 2

To determine whether *P. falciparum* infection influenced the frequency of mosquito sugar feeding, *An. coluzzii* females were exposed to infectious blood from a naturally-infected volunteer (parasite isolate D: 696 gametocytes/ul of blood). On 7 dpbm, the level of oocyst infection was measured by dissecting the midguts of 30 females. The proportion of infected females was 0.9 (27/30) with a mean number of 33 ± 5 oocysts. On the 13th day post infection, 30 females were placed in plastic cups with white filter paper at the bottom. The same day at 17:00 the cotton balls soaked with 10% glucose solution containing a blue dye (1 mg mL−1 Fast Green FCF/Xylene cyanole FF; Sigma-Aldrich) were placed on top of the cup gauze. The next day at 07:00, the colored cottons were removed and at 16:00 the presence or absence of mosquito droppings on the white filter paper were recorded. The same day at 17:00 this procedure was repeated with the same mosquitoes, but with a 10% glucose solution containing a yellow dye (1 mg mL−1 Acid Yellow 17; Sigma-Aldrich, St Louis, Missouri) (Ignell et al., 2010). The papers were changed every day and blue and yellow dyes were switched between days. The color of the fecal dots (blue, yellow or green, when two consecutive sugar meals were mixed up) were also specified each day at 16:00. The assay lasted 11 days until 24 dpbm. Females that died before 17 dpbm (i.e. from which less than 3 colorimetric measures could be performed were excluded). At the end of this experiment, we obtained 16 females surviving beyond 17 dpbm and from which were gathered 3 to 11 (median = 5) individual colorimetric measures. DNA present on cotton samples and carcasses of these 16 females was isolated and sporozoites quantified as in experiments 1 and 2.

#### 3B PSU

Fifteen *An. stephensi* were fed with colored sugar solution by providing 2 ml of sugar and dye added to medium size cotton wool pads (750 μl of blue food dye mixed with 15 ml 10% glucose solution). The cotton wool was held in place in the cap of plastic tube, glued to the bottom of a clear plastic cup, which was inverted over filter paper. The filter paper was held in place by a petri dish lid underneath to prevent mosquito escape from this cage and changed daily at 15:00 each day for one week. Mosquitoes were knocked down for 2.5 minutes in the freezer (−20 °C) to sort them randomly into cups. The 15 mosquitoes used in the experiment were previously blood fed at 6 days of age. These mosquitoes were used three days after their blood meal to encourage complete digestion and monitored daily thereafter. Color dots from fecal material from 15 *An. stephensi* individuals were counted on the daily filter paper samples.

A second experiment was conducted to test the effect of temperature on sugar feeding in two species as estimated by fecal dot production (a measure of digestion rate). A set of 45 *An. stephensi* and 45 mosquitoes of the G3 strain were treated the same way and set up at 9 days of age in inverted plastic cups as before. These were further divided into three temperature treatments with 15 mosquitoes per treatment and were housed at either 20 °C, 27 °C, and 32 °C and monitored for colored fecal dot production for eight days.Blue or green artificial food dye was used because pilot tests using 5 colors of various natural and artificial food dyes (as well as no dye) for sets of 5 *An. stephensi* mosquitoes housed and maintained in this way showed while there was no effect of any dye (compared to no dye) on mortality, the green and blue artificial food dyes were easier to see in fecal dots. Red dye resulted in fewer fecal dots (either from lack of feeding or a difference in processing) and yellow and natural red derived from beets were slightly more difficult to see.

### Experiment 4. Infection duration

Both at IRSS and at PSU, mosquitoes were tested for their ability to expel sporozoites on sugar cottons throughout their lives.

#### 4A IRSS

Infected females in individual tubes had cotton substrate collected every three days from 23 dpbm until the mosquitoes died. These “late infection” cottons (23-54 dpbm) were used for DNA extraction using the Qiagen technique before amplification using qPCR as described above.

#### 4B PSU

*An. stephensi* mosquitoes were kept in cups of either: a single mosquito, group of 4, group of 5, or a large group of more than 100, respectively and infected as described in 3B. Mosquitoes had sugar soaked cotton substrates placed the netting on top of the cups from 14:00 to 9:30 changed and collected daily from 12-40 dpbm, and analyzed intermittently out to 40 dpbm. At 40 dpbm all surviving mosquitoes were dissected and the terminal cottons from 40 dpbm were subjected to molecular analysis.

An additional 43 individually maintained infectious *An. stephensi* were sampled for the presence of parasites in the saliva every 5 days from 35 dpbm until death using sugar soaked 15mg cotton pads. Mortality was such that final sampling included 30dpbm, N=43; 35dpbm, N=36; 40dpbm, N=21; 45dpbm, N=11; and 50dpbm, N=3, at which point the experiment was terminated. No accompanying dissections were performed on these individuals, but from other data collected from mosquitoes fed under the same conditions, infection prevalence was nearly 100% and intensity of sporozoites was high at 15 days post infection. Extraction and detection of parasite DNA from both assays of the 4B experiment was done from the 15mg cotton samples as described above.

### Statistical analyses

All statistical analyses were performed using R (version 4.0.2). Logistic regression by generalized linear models (GLM, binomial errors, logit link, or quasibinomial errors) were used to test the effect of parasite isolates and day post blood meal (dpbm) on (i) the proportion of infected mosquitoes with ruptured oocysts (experiment 1), (ii) the proportion of ruptured oocysts (experiment 1), (iii) the proportion of oocyst-infected mosquitoes with sporozoites in their head and thorax (experiment 1). Since the outcome variable as part of the “spit” assay is the time when sporozoites are first detected in cottons from individual mosquito, we used the statistical approach specifically developed for investigating the time a specified event takes to happen. Cox proportional hazard models were therefore performed to test for statistical difference in sporozoite appearance time (or EIP by extension) among parasite isolates (experiment 1) and mosquito species (experiment 2). We also analyzed the relationship between the EIP and the lifespan of infected mosquitoes (experiment 2) using a generalized linear model (GLM). Model simplification used stepwise removal of terms, followed by likelihood ratio tests (LRT). Term removals that significantly reduced explanatory power (P < 0.05) were retained in the minimal adequate model (Crawley, 2012).

## Results

### Experiment 1: Comparing estimates of parasite’s EIP between the classic dissection approach and the non-destructive individual “spit” assay

#### 1.1 Destructive approach: mosquito dissection and microscopic observation

A total of 121 mosquito females exposed to parasite isolate A and 114 to isolate B were dissected from 8 to 16 dpbm (between 8 and 20 females / day, median = 14) to assess microscopically the presence and number of oocysts in the midguts and of sporozoites in salivary glands. Salivary gland infections were also confirmed through qPCR. The infection rate was high with 117/121 (96.7 %) and 114/114 (100%) of females exposed respectively to isolate A and B harboring parasite oocysts in their midguts (**Supplementary data 4,**Figure S4A). The gametocytemia of isolate B (1208 gam/ul) was higher than that of isolate A (168 gam/ul), resulting in strong difference in the number of developing oocysts between the two isolates (B: 191.65 ± 21, A: 13.86 ± 2, **Supplementary data 4,**Figure S4B, *LRT X^2^1* = 24.46, P < 0.001). Similar patterns of infection were observed in salivary glands for the sporozoite stages (**Supplementary data 4,**Figure S4 C and D).

The few uninfected mosquitoes (n= 4 individuals from isolate A) were excluded from the analysis of the EIP. None of the dissected mosquitoes at 8 and 9 dpbm exhibited ruptured oocysts and the first observations occurred at 10 dpbm for both isolates. As expected, there was a highly significant positive relationship between time post-bloodmeal and the proportion of mosquitoes with ruptured oocysts (*LRT X^2^1* = 147, P < 0.001, Figure 1A). The timing of rupturing was similar between the two parasite isolates (*LRT X^2^1* = 1.39, P = 0.24, Figure 1A). Using this metric, the estimated EIP_50_ from the binomial model was 9.98 days for isolate A and 9.49 for isolate B.

**Figure 1:**
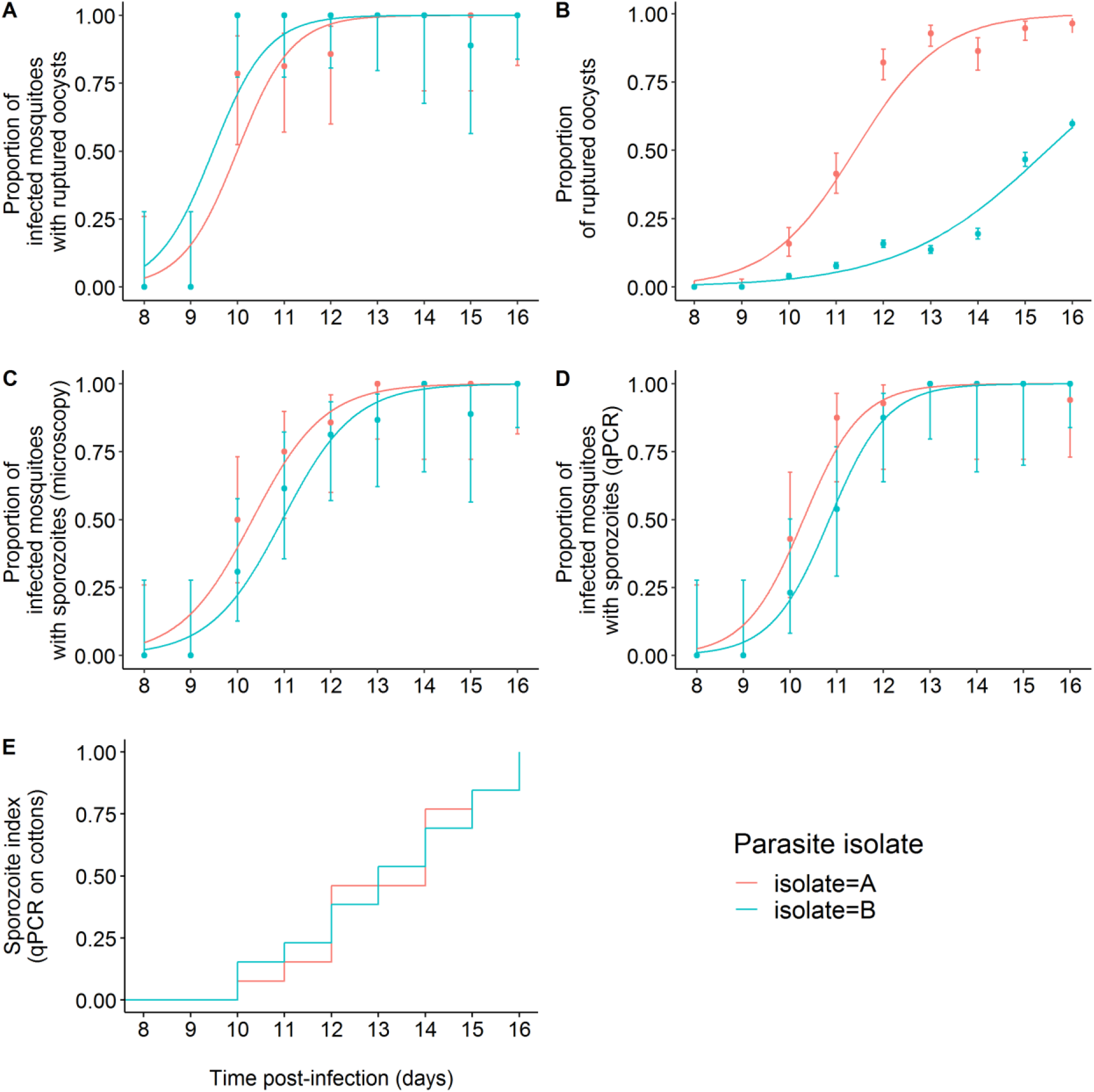
The extrinsic incubation period of *Plasmodium falciparum* estimated using classical dissection approaches (A-D) and a novel non-destructive assay (E). (A) Proportion of infected mosquitoes with ruptured oocysts (± 95% CI) from 8 to 16 dpbm, expressed as the number of mosquitoes with at least one ruptured oocyst out of the total number of infected mosquitoes (i.e. harboring either intact and/or ruptured oocysts) for two parasite isolates. The lines represent best-fit logistic growth curves for each isolate. (B) Proportion of ruptured oocysts (± 95% CI), expressed as the number of ruptured oocysts out of the total number of oocysts (intact + ruptured). The lines represent best-fit logistic growth curves for each isolate. (C) Proportion of oocyst-infected mosquitoes with microscope-identified sporozoites in the salivary glands (± 95% CI), expressed as the number of oocyst-infected mosquitoes harboring sporozoites in their salivary glands out of the total number of infected mosquitoes. The lines represent best-fit logistic growth curves for each isolate. (D) Same as (C) but the presence of sporozoite was detected using qPCR. A to D: Sample size = 8 to 20 midguts /day/isolate (median=14). (E) Kaplan-Meier curves representing the temporal dynamics of sporozoites appearance in small pieces of cotton used to collect saliva from individual mosquitoes.

The proportion of ruptured oocysts was higher in mosquitoes exposed to isolate A than B (A: 879 ruptured oocysts out of 1503 counted oocysts (58.5%) from 8 – 16 dpbm, B: 3 589 / 19 035 (18.9%) over the same period of time, *LRT X^2^*= 1045, P < 0.001, Figure 1B). This suggests a negative effect of density on parasite growth at the oocyst level, while no such density-dependent effect was observed on the proportion of mosquitoes with ruptured oocysts (Figure 1A) or with disseminated sporozoites (Figure 1C, D).

Finally, the proportion of mosquitoes with disseminated sporozoites in their salivary glands, the most epidemiologically-relevant metric, did not vary between parasite isolate, be it measured through microscopic observation (*LRT X^2^1* = 0.96, P = 0.33, Figure 1C) or qPCR (*LRT X^2^1* = 0.71, P=0.40, Figure 1D). The proportion of sporozoite-infested salivary glands increased with dpbm similarly for both parasite isolate (i.e. no significant dpbm by isolate interaction) and regardless of the method used. Using this metric, the estimated EIP_50_ from the binomial models were 10.35 (microscopy) and 10.26 days (qPCR) for isolate A and 10.93 (microscopy) and 10.85 days for isolate B.

The estimated time required for the sporozoites to migrate and invade the mosquito salivary glands following the egress from the oocysts was thus 9 hours for isolate A (i.e. EIP_50_ derived from the sporozoite observation in salivary glands (fig 1C) - EIP_50_ derived from the oocyst rupturing data (fig 1A): 10.35 days - 9.98 days), and 34 hours (10.93 days - 9.49 days) for isolate B.

#### 1.2 Non-destructive approach: the individual “spit” assay

Forty-two (42) females (21 per parasite isolate) were individually placed in tubes for saliva collection from 8 to 16 dpbm. Mosquito survival over the collection period is described in the **Supplementary data 4** (Figure S4E). At the end of the experiment on 16 dpbm, 37 females (17 in A and 20 in B) were identified as infected using qPCR. Of these, 26 (13 in each isolate) produced cottons containing detectable traces of parasite DNA by qPCR (i.e. positive cottons). The infected females that did not produce any positive cottons from 8 to 16 dpbm (4 females for isolate A and 7 for B) were excluded from the analysis because no EIP values can be derived from these samples. A total of 214 cottons (112 for A and 102 for B) collected from 26 females were thus analyzed.

Similar to microscopic observation, the first positive cottons occurred on day 10 for each isolate. The parasite EIP50 at 27°C using this assay was 14 days for isolate A and 13 days for isolate B (isolate effect: *LRT X^2^1* = 0.001, P = 0.97, Figure 1E).

### Experiment 2: Individual estimation of EIP in different mosquito species

#### 2A IRSS

The saliva of 20 *An. arabiensis*, 20 *An. coluzzii*, and 20 *An. gambiae* fed with the blood from a naturally infected gametocyte carrier (parasite isolate C) was collected using the “spit” assay from 8 to 20 dpbm. Of these, 18 *An. arabiensis*, 17 *An. coluzzii, and* 20 *An. gambiae* were confirmed as infected using qPCR on carcasses of dead mosquitoes. Cottons collected from uninfected females (n = 5) were discarded. Two infected females (one *An. gambiae* and one *An. coluzzii*) died at 7 dpbm before the collection of saliva has started (full survival results are given in **Supplementary data 5,**Figure S5.2). A total of 368 cotton samples from 53 females (18 *An. arabiensis*, and 16 *An. coluzzii*, 19 *An. gambiae*,) were therefore analyzed using qPCR. The proportion of positive cotton balls and the proportion of mosquitoes generating positive cotton samples are given in Table 1 for each species. Over the collection period from 8 to 20 dpbm, 19 individuals (2 *An. arabiensis*, 10 *An. coluzzii* and 7 *An. gambiae*), of the 53 infected females used to collect the saliva, never generated positive cotton samples (Table 1). This was mainly due to early mortality prior to the sporozoites invasion of mosquito salivary glands as this number fell to three (one from each mosquito species) between 13 to 20 dpbm.

**Table 1:** Proportion of mosquitoes producing at least one positive cotton over the collection period and proportion of positive cottons both over the collection period and after the first positive detection (“post-EIP”) for each anopheline species.

|  | collection period | *Anopheles arabiensis* | *Anopheles coluzzii* | *Anopheles gambiae* | *Anopheles stephensi* |
| --- | --- | --- | --- | --- | --- |
| Proportion of infected mosquitoes generating at least one positive cotton | 8-20 dpbm | 0.89 (16/18) | 0.38 (6/16) | 0.65 (12/19) | 0.93 (14/15) |
| Proportion of positive cottons | 8-20 dpbm | 0.26 (40/152) | 0.08 (7/93) | 0.33 (40/123) | 0.52 (99/190) |
| Proportion of positive cottons | post EIP | 0.69 (40/68) | 0.27 (7/26) | 0.71 (40/56) | 0.81 (99/122) |

EIP, defined as the time between the infectious blood meal and the first day of positive molecular detection by qPCR of *P. falciparum* from the cotton used to collect saliva, varied among species (LRT *X^2^*2= 8, P=0.018, Fig. 2A). The shortest EIP_50_ was observed in *An. gambiae* (11 days, min: 9, max: 16), followed by *An. coluzzii* (11.5 days, min: 9, max: 13) and *An. arabiensis* (13.5 days, min: 9, max: 18). Following parasite invasion of their salivary glands (i.e. the day when parasite DNA were first detected in cottons), individual females did not systematically generated positive cotton samples (Table 1). The “proportion of positive cottons post-EIP” refers to the sensitivity of our assay; that is, its ability to correctly detect parasite DNA in cotton samples collected on the days following the first day of positive detection. In other words, this is the proportion of cotton samples that tested positive for *P. falciparum* among those that were used to collect saliva of females that previously generated a positive cotton. Sensitivity was 0.69, 0.27, and 0.71 for *An. arabiensis, coluzzii,* and *gambiae* respectively (Table 1). Thus, *An. coluzzii* tended to deposit detectable quantity of sporozoites on cottons less frequently than the two other mosquito species following the parasite’s EIP.

**Figure 2:**
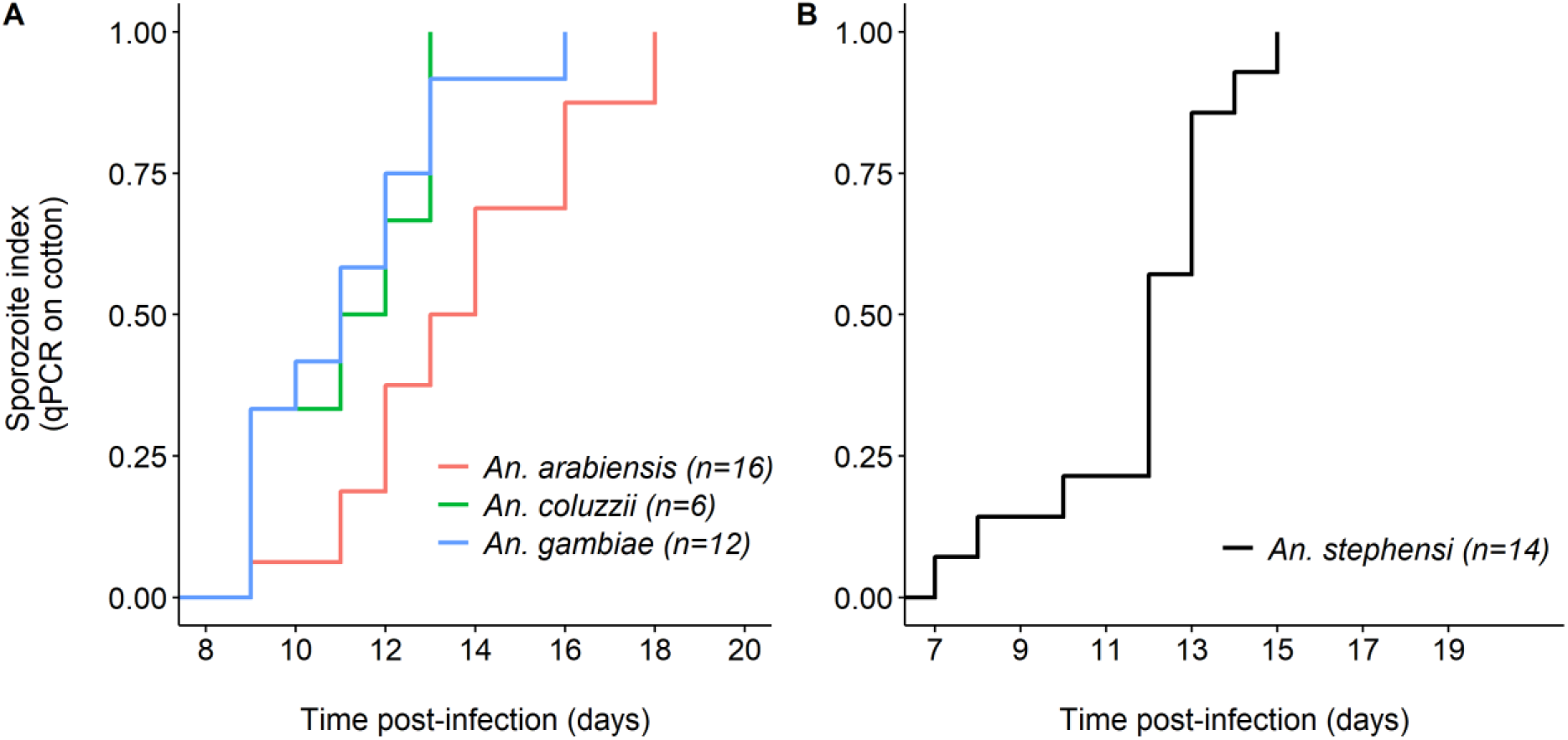
The extrinsic incubation period of *Plasmodium falciparum* in four Anopheles mosquito species. (A) Kaplan-Meier curves representing the temporal dynamics of sporozoite appearance in small pieces of cotton used to collect saliva from individual mosquitoes in the three major African vectors *An. arabiensis* (red), *An. gambiae* (blue) *and An. coluzzii* (green). (B) Same as (A) but for *An. stephensi*. The numbers in brackets indicate the number of females for each species of mosquito that generated at least one positive cotton.

A major driver of cotton positivity rate following parasite’s invasion of salivary gland was the infection intensity in females used to collect the saliva. First, *An. coluzzii* mosquitoes, the species with the lowest sensitivity (Table 1), also displayed lower infection intensity than *An. arabiensis* and *An. gambiae* **(Supplementary data 5,** figure S5.1A). Second, there was a positive relationship between the probability to generate Pf-positive cotton samples and infection intensity in individual females (LRT X^2^1=10, P=0.002, figure S5.1B), regardless of mosquito species (species by sporozoite intensity (Cp) interaction: LRT X^2^1=1.9, P=0.4, figure S5.1B). However, there was no relationship between the Cp of females used to collect saliva and the Cp of positive cottons (LRT X^2^1=1.9, P=0.4).

#### 2B PSU

Parasite-positive cotton samples were detected in *An. stephensi* (PSU). Of the 15 mosquitoes sampled, 14/15 *An. stephensi* individuals generated positive cotton samples at least once during the sampling period. After the dissection of all surviving mosquitoes at the end of the experiments, it was found that 12/12 *An. stephensi* were infected with salivary glands found harboring sporozoites. EIP50 similarly defined as in 2A was 12 days (Fig. 2B).

The spit assay makes it possible to study the links between different traits at the mosquito individual level. For the 2A IRSS experiment, cotton samples were collected daily up to 20 dpbm but the females used were kept in tubes until their death, thus making it possible to link the EIP with mosquito lifespan at the individual level. There was a significant positive correlation between EIP and mosquito longevity (LRT X^2^1= 21, P = 0.035, figure 3) such that short EIPs were also associated with short mosquito lifespan. There was no mosquito longevity by species interaction on EIP (LRT X^2^2=25, P=0.07, Figure 3), although 6 observations in *An. coluzzii* (very low statistical power) pointed to a negative correlation in this species.

**Figure 3:**
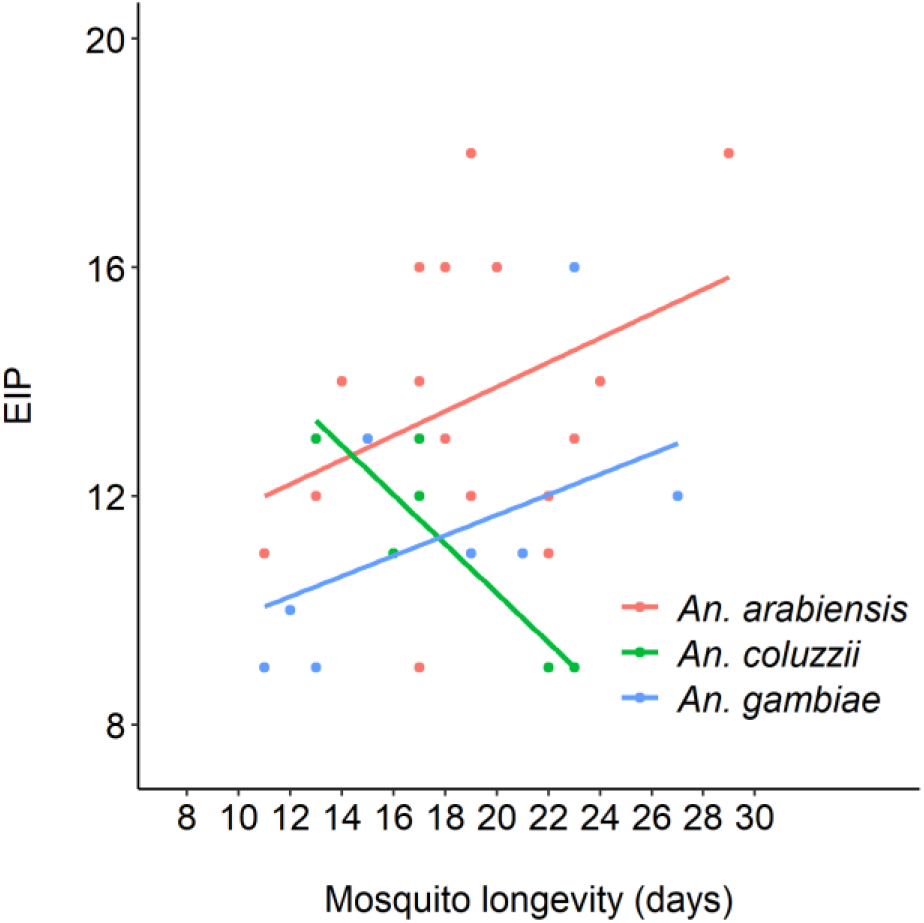
Relationship between the extrinsic incubation period (EIP) of *Plasmodium falciparum* and the lifespan of individual mosquitoes from three mosquito vector species.

### Experiment 3: Determination of mosquito sugar feeding rate

#### 3A IRSS

##### Assay 1

The overall mosquito survival rate during the 8 days-assay was 89%, with *An. gambiae* dying at a faster rate than both *An. arabiensis* and *An. coluzzii* (8/30 vs 0/30 and 3/30, *X^2^*2 = 11, P = 0.004). Dead mosquitoes were excluded from the analysis. A meal was scored as having been taken when filter papers at the bottom of the cups contained either blue, yellow or green fecal dots. The papers were changed daily and the presence of colored fecal dots was scored daily. Of the 79 mosquito survivors, 31 individuals generated colored filter papers all 8 collection days. The average number of positive days was 6.89 ± 0.14 (min: 2 days, median: 7 days, max: 8 days). The overall detected sugar feeding frequency over this period was 544 meals/632 feeding opportunities = 86%. The sugar feeding rate of *An. arabiensis* and *An. coluzzii* females was higher than that of *An. gambiae* (89 %, 91 % and 76 % respectively, GLMM with mosquito identity considered as a random effect: *X^2^*2= 13.9, P< 0.001).

##### Assay 2

The 16 females used to assess the presence of colored fecal dots from 14 to 24 dpbm were all infected (mean cp = 22.2 ± 1.37). A meal was scored as positive when filter papers at the bottom of the cups contained either blue, yellow or green fecal dots. A total of 91 colored cottons soaked in 10% glucose were retrieved from these infected females over the collection period, of which 63 were positive to *P. falciparum* (i.e. a 69% sensitivity) (Table 2, **Supplementary data 6**). The color of the cotton (yellow or blue) had no incidence on the probability to detect *P. falciparum* (*X^2^*1= 0.008, P=0.93). Colored dots were observed on 61 filter papers of a total of 91 observations; that is, a daily feeding frequency of 67 %.

**Table 2:**
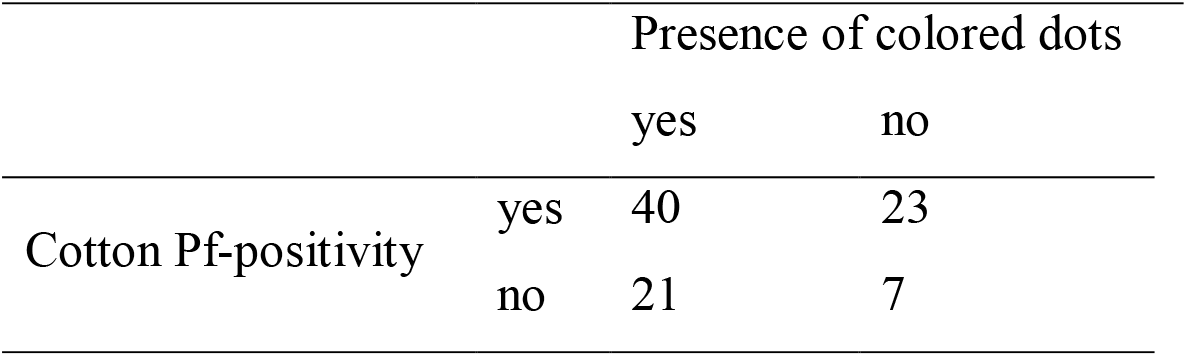
Association between sugar feeding (presence of colored dots) and production of Pf-positive cotton samples (cotton Pf-positivity).

Mosquitoes expectorating sporozoites of *P. falciparum* on cotton pads during the night (as evidenced by Pf-positive cotton samples, n=63) egested colored droppings on 40 occasions (63.4%). Contrary to our prediction, the absence of *P. falciparum* in cottons was not necessarily associated to the absence of colored dots: there were 21 occurrences of parasite-negative cotton samples but positive sugar meals, and only 7 occurrences of parasite-negative cotton samples and negative sugar meals (Table 2). Starting from dpbm15, there was a high proportion of green dots or combinations of green + yellow dots or green + blue dots. This shows that colored dots observed at 16:00 on a given day did not necessarily come from the digestion of a sugar meal taken during the same night, but could also result from previous sugar meals, meaning that mosquito sugar digestion can last > 24h. These results suggest that measuring colored fecal dots does not reliably provide evidence of either sugar feeding by night, or of transmission of parasites to cotton pads.

#### 3B PSU

The 15 *An. stephensi* mosquitoes maintained at 27 °C for 1 week produced sugar fecal dots most of days, suggesting a regular feeding. Due to some mortality, there were only 79 sample-days, but on 66 of these days mosquitoes produced at least 4 or more new dots on filter paper, suggesting they were likely feeding that day on the dyed sugar meal about 83.5% of days. No dots were observed on 8 sample days, or 10.1% of the time, with some variation of days with fewer than 4 dots observed making up the difference, where it was unclear if they were producing new dots or still digesting the previous day’s meal.

Color fecal dot production was monitored for *An. stephensi* (8 days) or G3 strain mosquitoes (7 days) housed at either 20 °C, 27 °C, and 32 °C. As might be expected, G3 strain mosquitoes produced fewer fecal dots at lower temperatures and greater numbers of dots at high temperatures. *An. stephensi* showed a similar pattern, although mortality at 32 °C was very high in this test which could have affected average dot production due to some individuals dying before digestion was completed (Table 3).

**Table 3.**
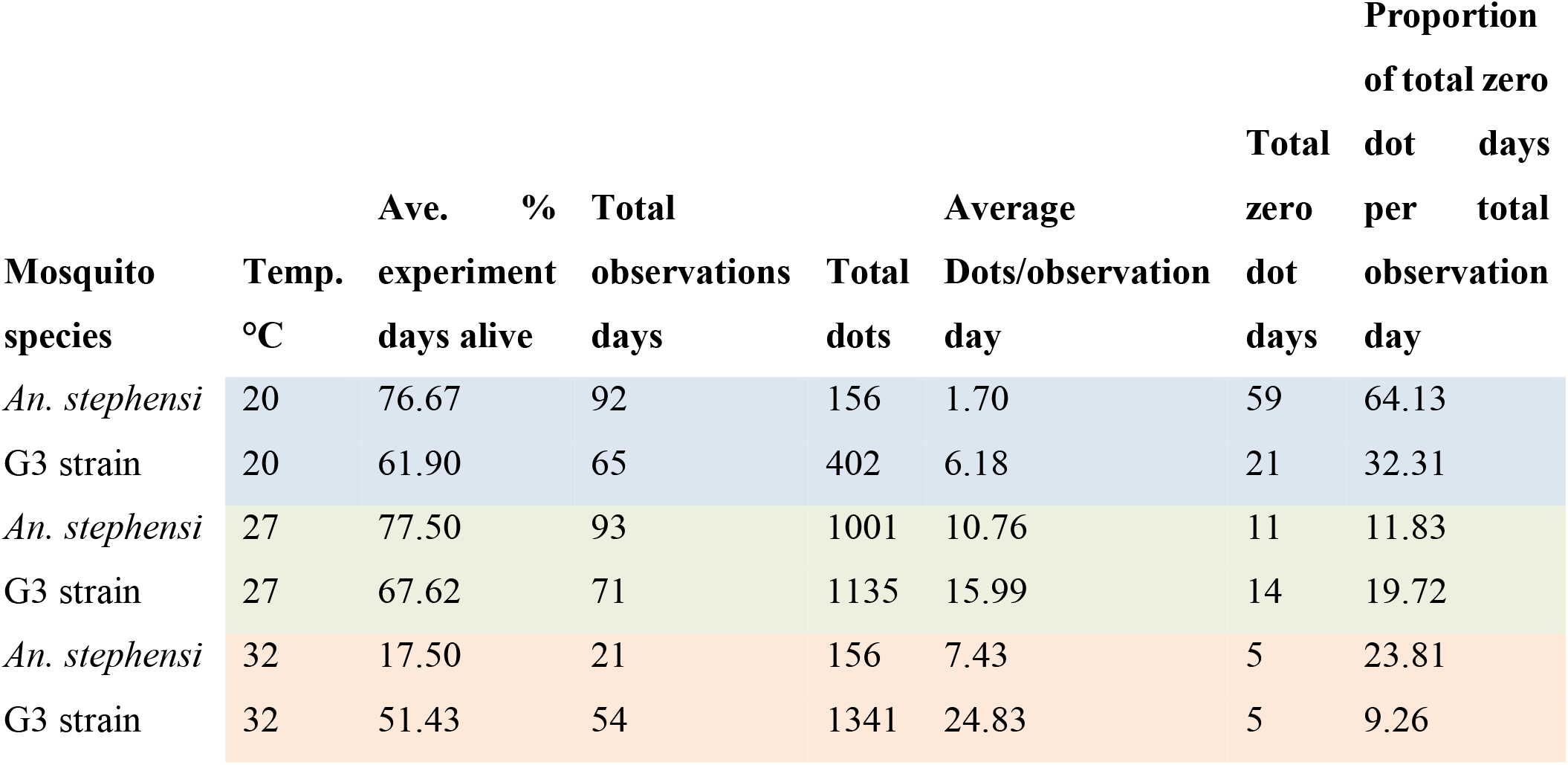
Summary of sugar feeding results comparing two species at three temperatures.

Generally, these results suggest that using the sugar-feeding assay to detect parasites will be more effective at warmer temperatures, and more false negatives are predicted at lower temperatures if mosquitoes are feeding less often. Around typical temperatures used in many infection studies (27 °C) mosquitoes are estimated to have fed about 80-90% of days sampled.

### Experiment 4. Infection duration

#### 4A IRSS

From a total of 34 cotton samples, 17 were positive. We could detect *P. falciparum* sporozoites in old cotton samples from 23, 26, 33, 36 and 39 dpbm.

#### 4B PSU

For samples at PSU tested in groups of 1,4,5, or 100+ in cups, surviving *An. stephensi* were carried through until day 40 dpbm, although only a few of these samples were analyzed. From this, there was evidence of salivary gland infection and transmission to cotton substrate as late as day 40 post infection from a group of 3 (those remaining from a set of 5), and also the larger group of 100, which by this point had about 20 mosquitoes remaining. Dissected mosquitoes were all found to have sporozoites in their salivary glands at 41 days post infection. This is evidence that infection can persist at least until day 40 post infection, and possibly until mosquito death.

## Discussion

Collectively, these data support that the assay we have developed is capable of monitoring individual mosquitoes repeatedly for infection status at transmission stages of malaria. We have further tested its robustness for use across various temperatures, finding it is functional ranging from 14 °C to 34 °C, and show that mosquitoes can remain infectious and continue to transmit parasites well into their old age, out to 40 days post-infection. The assay described here compared favorably to other assays with similar goals of passive monitoring (25). In another study, at 21 and 24 dpbm, respectively 31% and 55% of cotton wools collected from tubes hosting three *P*. *berghei-*infected females were positive (25). With mosquitoes kept individually, we found a detection rate (sensitivity) ranging from 0.27 to 0.81 depending on mosquito species (Table 1). The sensitivity of the presented assay was influenced by the amount of sporozoites in the salivary glands (Additional file 5). This may explain the observed inter-specific variation in sensitivity, with *An. coluzzii* displaying the lowest infection intensity and sensitivity (Additional file 5, S5.1A). To reach high level of sensitivity, we therefore recommend monitoring highly-infected females with this technique. The assay is easy and simple, only requiring basic molecular techniques of DNA extraction using a standard kit, and simple qPCR, a highly sensitive method for detecting the Cox1 gene in as few as six sporozoites (Marie et al., 2013). There has been much demand to determine whether a mosquito is transmitting sporozoites without dissecting it; including within human malaria research in areas as diverse as monitoring transmission in the field to developing better vaccines (Billingsley et al., 1991; Brugman et al., 2018; Churcher et al., 2017; Melanson et al., 2017). It can also be useful for animal models of malaria in the lab (Esperança et al., 2018). Additionally, it has the potential to advance studies on wild mosquito populations in the field on less-well understood species of avian malaria for example, where estimating infection prevalence in wild-caught mosquitoes when the timing of infection is typically unknown, or for generating experimental infections to examine host specificity in the lab if it is unclear if a mosquito is infectious since timing of EIP can vary by malaria species (Carlson et al., 2018). Applications of this low-tech innovation are many, and the most useful will be determined by those working in *Plasmodium* research and looking to avoid long hours of dissecting mosquito after mosquito to determine transmission, or through necessity if infected mosquitoes are limited. It would also open up the field to less-skilled/dexterous workers to aid in research as adding and removing sugar soaked cotton is non-technical, and extractions and qPCR routine compared to developing expertise at dissecting and scoring infections through microscopy.

Limitations of this non-destructive assay include a possible overestimation of the parasite’s EIP_50_ compared to classical measures using microscopic observations of salivary glands. In Experiment 1, the estimated EIP_50_ at 27°C of *P. falciparum* was 10 to 11 days using microscopy and 13 to 14 days with the non-destructive approach. Estimates derived from the second experiment using the non-destructive technique were shorter with EIP_50_ of 11 to 13 days depending on the vector species. The longer EIP estimates obtained with individual monitoring using the spit assay could be explained by the existence of false negative cotton samples. First, because of sensitivity < 100%, infectious females may deposit undetectable level of sporozoites onto cottons. This is especially true at the onset of salivary gland invasion when the amount of sporozoite is relatively low. Another source of false negatives is the biology of mosquito sugar feeding. Mosquitoes do not feed daily on sugar 100% of the time, and on average about 20% of potentially positive samples will be missed just because of skipped sugar feeds by infectious females within a 24 hour period, resulting in false negatives. This risk can be partially mitigated by feeding dyed sugar water and determining if mosquitoes are feeding at all by looking for colored fecal dots on the cage floor. If there are no dots for a few days, it is unlikely the mosquito is feeding on the cotton or that the sample will be positive for sporozoites (though digestion time will vary with temperature so the correlation is not perfect). Finally, sporozoites typically clump together and can be transmitted in groups in saliva (LI et al., 1992). This is further evidenced in observations by Frischknecht and collaborators (Frischknecht et al., 2004) where sporozoites were observed to move in clumps into salivary ducts for transmission. This suggests that transmission can be spotty and unpredictable by nature, even in infectious mosquitoes, yet this may be adaptive for the parasite if it allows the sporozoites to remain below a detection threshold in the vertebrate host upon inoculation, yet have enough transmitted at a time to establish infection (Aleshnick et al., 2020; Churcher et al., 2017; LI et al., 1992). It is also worth mentioning that to avoid wasting time on a feed that was not infectious, it is always best to check at least a few mosquitoes for oocysts and/or sporozoites using dissection at appropriate time points to ensure the infection was successful.

Longer EIPs values obtained using the spit assay compared to microscopic observations (experiment1) could also result from the existence of a delay between the sporozoite invasion of salivary gland (the trait measured by microscopic observations) and the expulsion of sporozoite in saliva (the trait measured by the spit assay). For example, in *Aedes aegypti* infected with *Plasmodium gallinaceum*, only about 80% of infectious mosquitoes released sporozoites during a forced salivation assay, and it was observed that there was a one-day delay between the appearance of parasites in the salivary glands and the sporozoites being released with saliva (Spielman and Rossignol, 1985). The difference between EIP values derived from microscopic observations and that from the detection in cotton samples therefore supports the existence of such a delay in *P. falciparum*-infected *Anopheles* mosquitoes. This experiment also showed that the first ruptured oocysts occurred at 9-10 dpbm and that the average time between oocyst rupture and the first invasions of salivary glands was 9 to 34 hours depending on the parasite isolate. The rate of oocyst rupturing was recently considered as the major driver of EIP and hence a key epidemiological parameter (Childs and Prosper, 2020). Our observations suggest that the day of first oocyst rupture +1 may then roughly indicate the time of salivary gland invasion. Moreover, we confirm that only a fraction of oocysts ruptures to release sporozoite and that this fraction seems to depend strongly on the oocyst intensity in the gut (figure 1B). Further work is needed to better understand variations in the delays between both the oocyst rupturing and salivary gland invasion and between invasion and injection with saliva.

Mosquito survivorship with the assay was high and long-term individual tracking revealed that sporozoite transfer onto cotton wools can persist at least until day 40 post-infection. Given the relatively short lifespan of *Anopheles* mosquitoes in field conditions (15-20 days) (Matthews et al., 2020) this result therefore suggests that mosquitoes likely remain infectious for life.

Perhaps the greatest utility of the assay is that it can be used for individual mosquitoes. Currently, nearly all estimates of sporozoite prevalence, or even estimates of EIP, are measured as averages of groups of mosquitoes being sampled. Even taking into account the possibility for false negatives using the assay, there are great opportunities to monitor aspects of effects of malaria infection on mosquitoes that we have not previously had the chance to explore. This could open up areas of research to look at parasite transmission over lifespan, individual variability in earliest transmission, possibilities to examine how fecundity is (or is not) affected by infection, and associations between transmission traits such as e.g. EIP, infection level, survival and fecundity. As one example, results from experiment 2 indicate a positive relationship between mosquito lifespan and the parasite’s EIP, suggesting the existence of a trade-off between two important transmission traits. A similar pattern was previously observed with a non-destructive sugar-feeding assay in the dengue – *Aedes aegypti* association (Ye et al., 2016). The trade-off between EIP and mosquito longevity could be mediated through parasite factors (e.g. more virulent parasite isolates also develop faster) and/or mosquito factors (e.g. poorly vigorous mosquitoes with low lifespan perspective are also more permissive to the development of the parasite). Additional work will be required to confirm this relationship. The non-destructive sugar-feeding assay developed here is a unique opportunity to quantify concomitantly multiple parasite and mosquito traits at the individual level and hence will contribute to a better understanding of the evolution of key epidemiological traits.

## Author Contributions

EG, JLW, MJ, and AB designed and tested molecular aspects of the assay, ES assisted in infectious feeds conducted Penn State and shared some infected mosquitoes and comparative data from his experiments. EG, TL, MBT and JLW devised the assay. EG, TL and JLW conducted experiments, and drafted the manuscript. BKY and EG performed the qPCR at IRSS. JLW did qPCR at Pennsylvania State University in Pennsylvania USA. EG and ND produced mosquitoes and performed experiments. EG, JLW and TL analysed the data. DFD, DFdSH, RSY, AGO, KRD and AC contributed to reagents/materials/analysis tools.

## Acknowledgements

Thanks to Deonna Soergel for help with dissections, Janet Teeple and Natalie Gahm for mosquito colony rearing assistance, and the Thomas and Read groups, Johanna Ohm, and Elizabeth McGraw for helpful discussion. Thanks to Fhallon Ware-Gilmore for sharing some infected mosquitoes. We thank gametocyte carrier volunteers for participating in this study and the local authorities for their support. Thanks to the infectivity group at the IRSS Bobo Dioulasso for technical support and discussions. MBT, JLW and ES were part-funded by NIH NIAID grant # R01AI110793 and National Science Foundation Ecology and Evolution of Infectious Diseases grant (DEB-1518681). The funders had no role in study design, data collection and analysis, decision to publish, or preparation of the manuscript.

## Supplementary Data

**Supplementary Data S1:** Text S1. Assay development and optimization. Figure S1.1. Experimental design. Table S1.1. Number of infected according to sample size and mosquito density. Figure S1.2. Proportion of sporozoite infection according to sample type. Different letters (a, b, c) indicate differences between infection status. Table S1.2. Number of infected according to sample type. Figure S1.3. Proportion of sporozoite infection according to sample type and parasite DNA extraction type. Table S1.3. Comparing effects of sample collection method and sample exposure time on sporozoite detection. Table S1.4. Evidence that our probe-based qPCR assay is functional, that sporozoites are deposited on sugar soaked filter paper squares, and that the assay works across mosquito species. Figure S1.4. qPCR for 12 samples from cups of mosquitoes with good sporozoite infection (high prevalence and intensity). Table S1.5. Bass Assay vs. SYBR assay comparison. Table S1.6. Samples: sporozoite standard mix from infectious *An. stephensi*, quantified 135,000 spz/ml using hemocytometer. 40 glands in 1.2 mls PBS, portioned as follows into either an Eppendorf tube (liquid samples), or onto 15 mg cotton treated either with sugar solution, or left dry “plain”. Samples were either frozen immediately “frozen” or left in the 27 °C, 80% rH conditions for 24 hrs prior to extraction and qPCR SYBR assay.

**Supplementary data S2:** Text S2. Testing the assay at various temperatures. Table S2.1. Comparing likelihood of sporozoite detection using this assay at various temperatures.

**Supplementary data S3:** Text S3. Oocyst rupturing and release of sporozoites. Figure S3.1. Immature developing oocysts. Figure S3.2. Mature and immature oocysts. Figure S3.3. Ruptured and unruptured mature oocysts. Figure S3.4. Ruptured oocysts.

**Supplementary data S4:** Text S4. Infection level and mosquito survival in experiment 1. (A) Oocyst prevalence (± 95% CI) on day 8-9 post-blood meal (dpbm). (B) Oocyst density at 8-9 dpbm. (C) Sporozoite prevalence (± 95% CI) at 14-16 dpbm. (D) Sporozoite density at 14-16 dpbm. (E) Survival of mosquito females used to collect saliva for each parasite isolates.

**Supplementary data S5:** Text S5. Infection level and mosquito survival in experiment 2. Figure S5.1A. Infection load in mosquito females used to collect saliva. Figure S5.1B. Estimated probability of *Plasmodium falciparum* carriage in cottons as a function of infection load in females used depositing saliva on these cottons. Figure S5.2. Survival of mosquito females used to collect saliva for each anopheline species.

**Supplementary data S6:** Text S6. Relationship between mosquito sugar feeding and *P. falciparum* positivity in cottons. Table S1: Evaluation of the presence colored fecal dots from 14 to 24 days after the infectious blood meal (dpbm) from 16 females infected with *P. falciparum*.

## Notes

### Competing Interest Statement

The authors have declared no competing interest.

