## Supplementary material for "Using a non-destructive sugar-feeding assay for sporozoite detection and estimating the extrinsic incubation period of *Plasmodium falciparum* in mosquito vectors": Supplementary data 1.pdf

**Supplementary data 1: Assay development and optimization**

Pilot experiments were conducted to determine the optimal substrate, extraction method and manipulation of substrate, assay duration, and minimum mosquito number in the sugar-feeding assay. Various detection methods using qPCR were also tested.

**METHODS**

**1. Optimizing mosquito sugar feeding substrate, manipulations, and mosquito density**

**Methods IRSS:** Laboratory-reared *An. gambiae* (Burkina) females were infected with two parasite isolates. At 7 days post blood meal, oocysts of some mosquito were examined to determine infection prevalence. At this stage, mosquitoes were placed individually or as groups of 2 or 3 placed in plastic drosophila tubes (diameter of 25 mm and height of 95 mm) covered on the top with a mosquito net (Figure S1-1) to collect mosquito saliva. Cotton balls and whatman papers soaked with 10% glucose solution were deposited on the top of the tubes for mosquitoes feeding. Several exposure times of cotton balls and whatman papers (~1cm x 0.5cm) for mosquito were used: 2 hours, 13 hours, 24 hours and 48 hours.
Following this exposition period, the substrates were placed individually in sterile 1.5 ml tubes and stored at -20 °C for further processing. The presence of sporozoite in cotton balls, in whatman filter paper and in the mosquitoes used to collect saliva (carcasses) was determined using one of three DNA extraction methods: CTAB, DNAzol and Qiagen Dneasy® Blood and Tissue Kits.

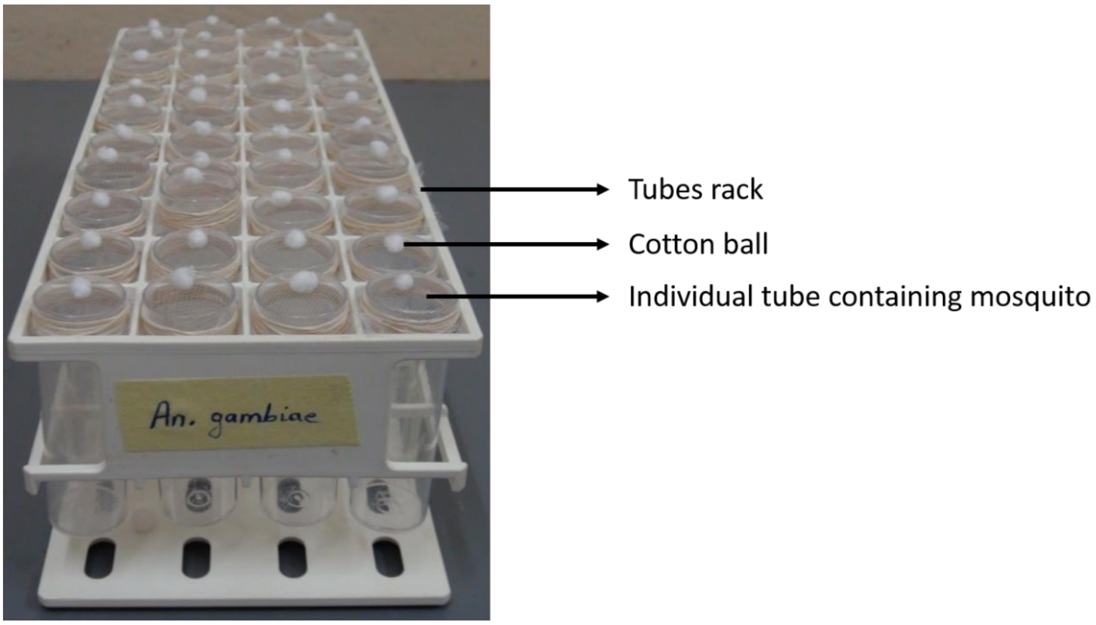


**Fig. S1.1: Experimental design**

**Methods PSU:** Mosquitoes (colony G3, a mix of the *An.gambiae* and *An. coluzzi* species versus *An. stephensi*) were housed in six 480 ml wax-lined paper cups total in groups of 120-150 and fed an infectious blood meal from *P. falciparum* culture, with 4 cups housing *An. stephensi* and 2 housing colony G3 mosquitoes. Cups were labelled according to mosquito species, and half of each set of cups was fed either dilution 1 (D1) or dilution 2 (D2) of the parasite culture. Dilution 1 had a higher concentration of parasite culture that resulted in higher prevalence and intensity of infection overall in mosquitoes compared to D2. All mosquitoes were maintained at 27 °C until day 16 post infection with both a large sugar soaked cotton pad with PABA. Mosquitoes were removed for dissection daily to provide oocyst intensity and prevalence and sporozoite prevalence data. On days 15 and 16 post infection, the approximately 20-50 remaining groups of mosquitoes in these 6 cups were provided a small 1 cm x 0.5 cm Whatman filter paper rectangle (~1cm x 0.5cm) with a 15 mg piece of 10% glucose sugar soaked cotton wool on top of the filter paper to wet it for sampling sporozoites added in addition to a larger sugar soaked cotton wool used for maintenance. Filter papers with cottons were left on the netting on the top of each cup for a 24 hour period and then was collected into -20 °C. Only the filter paper was used for DNA extraction by the Qiagen kit method, cotton discarded assuming mosquitoes could not probe through the paper. Results are described in the results section 3: **Alternative qPCR probe-based method and other qPCR comparisons** since these samples were analyzed using an alternative probe-based qPCR method.

Note (PSU): Other substrates were also tested, but sugar soaked cotton resulted in the best feeding success and lowest mosquito mortality, and was easiest to manipulate. Liquid assays were tried with uninfected *An. stephensi* mosquitoes based on (Ye et al., 2016), with either open caps filled with dyed sugar solution on the bottom of the cup, or sugar solution in tubes covered with thin parafilm made accessible from the top of the cup. *Anopheles* mosquitoes preferred to sugar feed from the top of the cage rather than on the bottom from a liquid pool. Moreover, extraction of a leftover blood provided in a meal from a membrane feeder to infectious mosquitoes also resulted in a sporozoite positive DNA sample, but was messy and labor intensive, and feeding compliance daily may be low (although many mosquitoes will blood feed daily if provided the opportunity). Other sugar meal substrates including grape gelatin cubes on or off of filter paper on top of the cage netting, or sugar cubes or pieces of sugar cubes either did allow mosquitoes to feed well, left sticky reside, molded, or couldn't be easily collected (pers. obs. JLW, data not shown).

### 2. DNA extraction methods comparison

Extraction methods tested included CTAB, DNAzol extraction, and extraction using a Qiagen DNeasy Blood and Tissue kit. For the Qiagen kit we also tested how long to leave the cotton in the tube during the extraction, and found the best results when it was carried through to the end of the extraction, with AE elution buffer placed on the cotton and spin at the last step.

**CTAB extraction**: Parasite DNA from cottons and Whatman filter papers was extracted with Cetyl Trimethyl Ammonium Bromide 2% (CTAB 2%). Each sample was ground in 200 µl of a 2% solution of CTAB, which allows the inactivation of cellular nucleases. Cellular lysis was accentuated by placing the crushed mosquitoes for 5 min in a water bath at 65 °C. Then, 200 µl of chloroform was added. Samples were mixed by inversion and then centrifuged for 5 minutes at 12,000 rotations per min (rpm). The supernatant was removed into another 1.5 ml Eppendorf tube. In the tubes containing the supernatant, 200 µl of isopropanol was added and samples were mixed by inversion. These samples were centrifuged for 15 min at 12,000 rpm. After centrifugation, the supernatant iso-propanol mixture was emptied. Then, 200 µl of 70% ethanol was added and tubes were centrifuged for 5 min at 12,000 rpm. After this, 70% ethanol was emptied from the tube and the pellet dried for 5 min maximum at speed-vac. Finally, the DNA was recovered in 20 µl of ultrapure water and left in suspension on the bench all night before being stored at -20°C (Doyle and Doyle, 1991). It was the same procedure for the extraction of parasite DNA from mosquito carcasses (head/thoraces).

**DNAzol extraction**: The samples of cotton and Whatman cards stored in 200 µl of DNAzol were ground. They were mixed by inversion and allowed to stand at room temperature for 5-10 min. At the end of this time, the samples were centrifuged at 14,000 rpm for 10 min. The supernatant was then transferred to a new 1.5 ml Eppendorf tube. The DNA was then precipitated by adding 200 µl of 100% ice-cold ethanol. The samples were mixed by inversion and incubated at -20 °C for 3 hours. Centrifugation of 14,000 rpm for 10 min was performed and the ethanol was removed. The pellet thus obtained was washed first with 75% ethanol at 14,000 rpm for 5 min and second with 75% ethanol at 7,500 rpm for 5 min. The ethanol was then removed by inversion and the pellet dried at Speed-Vac for 10 minutes. The DNA was dissolved in 50 µl of ultrapure water for half a day at room temperature (Ausubel et al., 2003). The same procedure was used for the extraction of parasite DNA from mosquito carcasses (head/thoraces).

**Qiagen DNeasy Blood and Tissue kit extraction**: 180 µl of ATL buffer are added in the 1.5 ml tubes containing the cotton balls or Whatman cards and then samples were crushed. 20 µl of proteinase K was added and then vortexed for 30 sec before being incubated at 56 °C for 10 minutes in a water bath for lysis. 200 µl of Buffer AL was first added to the samples and then 200 µl of absolute ethanol (100%). The samples were then vortexed for 30 sec. The mixture obtained was transferred (together with the cotton balls or Whatman filter paper) into the columns (DNeasy Mini Spin Column). The columns were centrifuged at 8,000 rpm for 1 min and then transferred to new Qiagen tubes. The previous tubes containing the flow-through liquid were discarded. 500 µl of AW1 buffer was added and the columns were then centrifuged at 8,000 rpm for 1 min for the first wash. The second set of tubes was then discarded and the columns transferred to a third set of Qiagen tubes. 500 µl of AW2 buffer was then added before centrifuging the columns at 14,000 rpm for 3 minutes for the second wash. The third set of tubes was then discarded and the columns were then gently removed so that they would not come into contact with the liquid contained in the tube. The columns were transferred into Eppendorf 1.5 ml sterile tubes. The DNA contained in the column was solubilized in 50 µl of AE buffer and the tubes were then centrifuged at 8,000 rpm for 1 minute. The columns were then removed and eluted liquid contained in the 1.5 ml sterile Eppendorf tubes was stored at -20 °C for subsequent PCR. It was the same procedure for the extraction of parasite DNA from mosquito carcasses (head/thoraces).

Extraction methods using the Qiagen DNeasy kit at PSU were identical with the following minor exceptions. Samples were incubated at 56 °C on a shaker table heat block for microcentrifuge tubes set at 1,200 rpm for at least 10 minutes rather than in a Bain-Marie for lysis. Samples were additionally vortexed for 30s with the addition of Buffer AL. DNA samples were stored at -80°C for later PCR rather than -20°C.

Additionally, in Pennsylvania, the Qiagen DNeasy kit was used to compare the extraction efficiency when the cotton substrate was removed at different time points during the extraction procedure.

### 3. Alternative qPCR probe-based method and other qPCR comparisons

In addition to the qPCR methods described in the main manuscript, additional primer and amplification methods were tested. None performed as well as the SYBR qPCR methods described in the main manuscript, this probe based method also worked, but with a higher detection threshold.

#### 3A: Probe-based qPCR

Parasites were quantified by quantitative real-time PCR by amplifying the cytochrome b gene of the *P. falciparum* parasite. 10 µl reaction conditions consisted of: 5 µl master mix, 1.2 µl forward primer, 1.2 µl reverse primer, 1.6 µl water, and 1 µl template DNA from the extracted product per reaction. The PCR kit PerfeCTa® qPCR FastMix®, UNG, Low ROX™, Cat #95078 from Quanta BioSciences (VWR) was used. The following primer sequences were developed and used: forward primer qPCR-PfF (TTA CAT CAG GAA TGT TAT TGC), reverse primer qPCR-PfR (ATA TTG GAT CTC CTG CAA AT). Final primer concentrations were 240 nM. Samples were run with a minimum of a water negative control and positive control of *P. falciparum* DNA derived from 2 µl of *P. falciparum* in tissue culture extracted with a substrate to mimic initial trials for mosquito sugar feeding. Sporozoites were quantified using a 7500 Fast Real-time PCR System instrument, with thermocycler conditions set for 20 seconds of activation at 95°C, denaturation for 3 seconds at 95°C, and 30 seconds at 60°C to anneal and extend the product for 40 cycles.

**3B: Bass assay and SYBR assay qPCR comparison**

We compared the SYBR assay we used most consistently as described in the main methods section in the manuscript (under qPCR methods for *P. falciparum*) to the probe-based Bass assay described in (Melanson et al., 2017) for detection of *P. falciparum*. A standard curve was created using 100 ul samples of a serially diluted sample of PBS spiked with *P. falciparum* NF54 strain culture placed onto 15mg cotton followed by extraction and comparative qPCR assays. Neat refers to the undiluted sample, -1 is diluted 1:10, -2 is diluted 1:100, -3 is diluted 1:1000, -4 is diluted 1:10,000. The unknowns tested in this assay are from sugar-soaked 15mg cotton substrate fed on overnight by *P. falciparum* infected *An. stephensi* mosquitoes on the $13^{th}$ day post infection.

**4. Generating a standard curve and testing how sample handling processes might affect sample loss or degradation:**

*Standard curve generation*

Sporozoite infected from salivary glands of 40 infected mosquitoes and collected into 1,200 ul of PBS, and homogenized. The number of sporozoites was quantified using a hemocytometer. 100ul of this liquid solution was used to generate a standard curve by extraction using the Qiagen DNeasy kit and eluted into 50 ul AE buffer (as all cotton samples were treated in the main experiment). The extracted DNA of the infected salivary gland homogenate was serially diluted 1:2 a total of 8 times (1:1 through 1:128) to generate dilutions for use in a standard curve. These were run in triplicate. Additionally 1 and 2 ul of extracted DNA template was used to compare the effect of DNA template concentration.

*Effects of heat, time, and processing on degradation*

The salivary gland homogenate was also used to test sample handling and measure degradation or loss in the sampling process. Different amounts (5 ul, 20 ul, or 100 ul) of PBS-sporozoite mixture were pipetted either a) as liquid into a new 1.5 ml Eppendorf tube b) onto 15 mg dry cotton, c) onto cotton that had been dipped in sugar and left overnight in 27°C to simulate the moist cotton sample normally collected in this assay. The samples in the b and c treatments were duplicated, and half were placed directly in the freezer, and half were left in for 24hrs at 27 °C 80% rH before being capped and frozen the next day. The 100 µl sample was noticeably oversaturated compared to typical cottons used in the assay, but moisture in the 5µl and 20µl samples seemed within the normal moisture range of experimental cotton collected. All samples were extracted in the same extraction round and qPCR'd the following day. qPCR samples were run in duplicate.

**RESULTS**

**<u>1. Optimizing mosquito sugar feeding substrate, manipulations, and mosquito density</u>**

**Results IRSS:**

*Mosquito number for assay*

Mosquitoes housed in groups of 3 appear to give better sporozoite detection results than those housed individually or in groups of 2. However, since the test also worked for mosquitoes kept individually, and since individual conditioning is the only way to obtain a reliable estimate of the EIP or other relevant characteristics of the mosquito's life cycle, we chose use mosquito single conditioning for future experiments.

**Table S1.1: Number of infected according to sample size and mosquito density.**
The detection rate expressed the proportion of *P. falciparum*-positive cotton or Whatman filter paper detected in qPCR on the number of samples analyzed in qPCR (according to sample type).

| **Mosquito density** | **Sample** | **Infected (number)** | **Total (number)** | **Detection rate from 8 to 41 dpbm (%)** | **Chi-test** |
|---|---|---|---|---|---|
| Individual | Cotton | 73 | 213 | 34 | $X^2_1$=5.41; P=0.02 |
| | Whatman filter paper | 40 | 171 | 23 | |
| 2 | Cotton | 19 | 53 | 36 | $X^2_1$=2.02; P=0.15 |
| | Whatman filter paper | 11 | 48 | 23 | |
| 3 | Cotton | 31 | 61 | 51 | $X^2_1$=2.70; P=0.10 |
| | Whatman filter paper | 22 | 61 | 36 | |

*Optimal substrate*

**IRSS:** The sporozoites detection was higher in cotton balls compared to Whatman filter paper. **(Table S1.1&S1.2&S1.3, Figs S1.2&S1.3).** Sporozoites was successfully detected in 123 of 328 cotton balls samples (37.5%) and in 73 of 280 Whatman filter papers samples (26%), ($X^2_2$=84.8; P=0.002; Fig. 1; Table S2). The detection rate in cottons and in papers was low compared to the infection rate of sporozoites in females (57/57=100%) used to collect these cottons balls and whatman papers ($X^2_2$=96.7; P<0.0001; Fig. S1.2; Table S1.2).

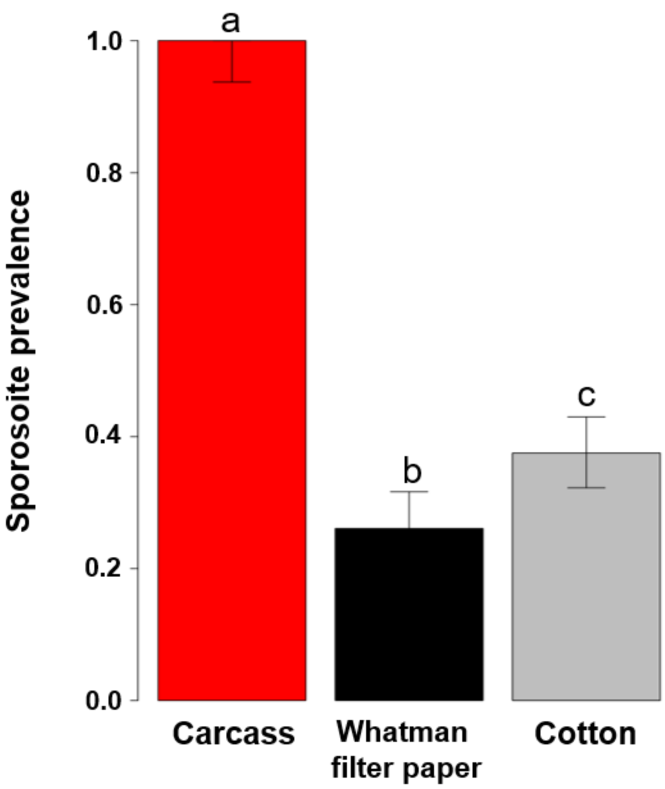


Fig. S1.2: Proportion of sporozoite infection according to sample type. Different letters (a, b, c) indicate differences between infection status. Red bar: carcasses of females used to collect saliva, black bar: whatman filter paper and gray bar: cotton.

**Table S1.2: Number of infected according to sample type**

| **Sample** | **Infected** | **Non infected** | **Infected Proportion** | **Chi-test** |
|---|---|---|---|---|
| Cotton | 123 | 205 | 37% | $X^2_2$=9.03; P<0.002 |
| Whatman filter paper | 73 | 207 | 26% | |
| Carcass | 57 | 57 | 100% | $X^2_2$=109.89; P<0.0001 |

### <u>2. Optimal extraction method</u>

The Qiagen extraction technique gives a better sporozoite detection rate than CTAB and DNAzol. In addition, for all three extraction techniques, mosquitoes produced more positive cotton balls (Qiagen extraction) than positive Whatman filter paper ($X^2_1$=15.36; P<0.0001; Fig.S1.3).

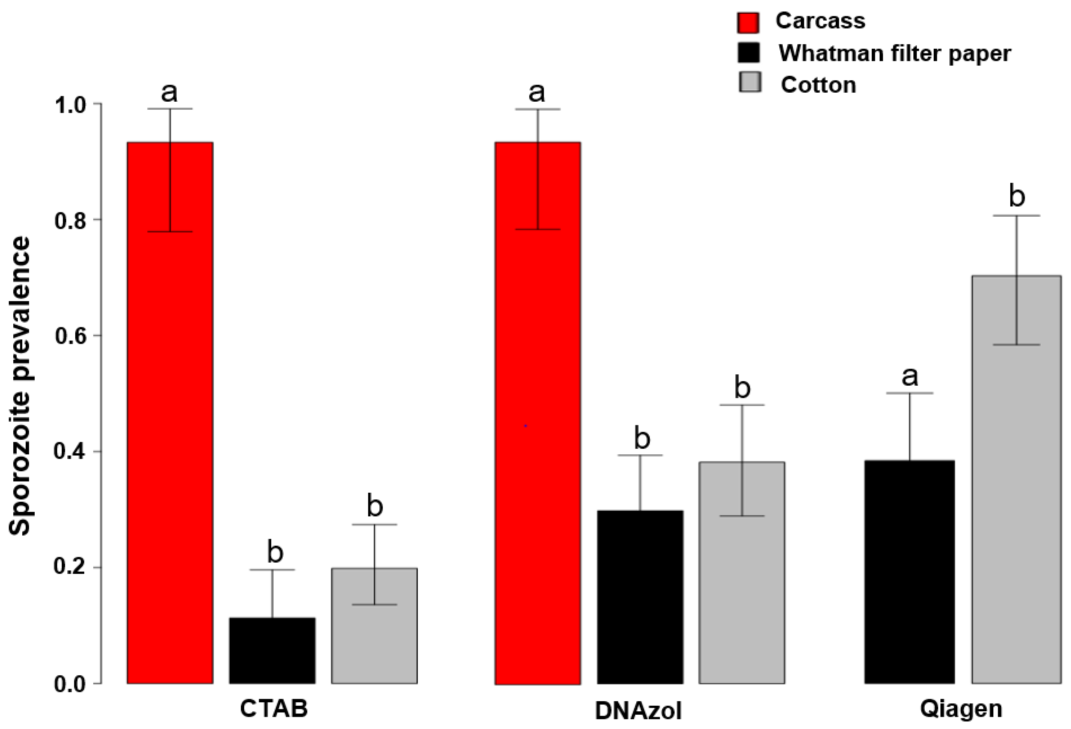


**Fig. S1.3**: Proportion of sporozoite detection according to sample type and parasite DNA extraction methods. Different letters (a, b) indicate differences in parasite detection between substrates compared within an extraction treatment. Red bar: carcasses of females used to collect saliva, black bar: whatman filter paper and gray bar: cotton. Note: no Qiagen-extracted carcasses

*Exposure time / duration of assay*

To compare exposure time, samples were divided into two groups. Exposure time in the first group was set at ≤ 13 hrs and included both cotton samples placed on gauze of infected mosquito housing for 13 hrs and Whatman filter papers similarly placed for 2 hrs. Exposure time in the second group was set to ≥ 24 hrs to increase sample size (24 and 48 hrs). More samples were detected positive using the exposure time of ≤ 13. Within this group, the more positive samples were detected using cotton compared to papers ($X^2_1$=10.68; P=0.001, Table S3).

**Table S1.3: Comparing effects of sample collection method and sample exposure time on sporozoite detection**

| Exposition time (hour) | Sample | Infected (number) | Total (number) | Infection proportion (%) | Chi-test |
|---|---|---|---|---|---|
| ≤ 13 | Cotton | 95 | 230 | 41 | $X^2_1$=10,68; P=0,001 |
| | Whatman filter paper | 52 | 198 | 26 | |
| ≥ 24 | Cotton | 28 | 98 | 28 | $X^2_1$=19,77; P=0,66 |
| | Whatman filter paper | 21 | 82 | 26 | |

Increasing the exposure time of cotton balls or Whatman filter paper did not improve sporozoite detection. A single exposure period (13 hours) was sufficient.

### 3. Alternative qPCR probe-based method and other qPCR comparisons

**3A: Probe based qPCR assay:** Cp values from MJ (Matthew Jones) method of primer/probe qPCRs (lower than 35 cycles is considered positive), paired with sporozoite prevalence data from dissections. Ordering of sample identification (ID) is researcher, dilution (D1 is more concentrated than D2), Anopheline mosquito species, and number of days post infectious feed on which the sample of filter paper with sugar-soaked cotton on top was collected. There were 6 cups used (4 *An. stephensi*, 2 *An. gambiae/coluzzi* G3 strain) of around 20-50 mosquitoes for the previous 24 hours as described in methods section S.1B. Only the filter paper was further processed (small sugared cottons on top of filter papers were discarded).

**Table S1.4. Evidence that our probe-based qPCR assay is functional, that sporozoites are deposited on sugar soaked filter paper squares, and that the assay works across mosquito species.**

| Sample ID | Cp value | Oocyst prevalence and (intensity) 8dpi by dissection | Spz prev. (closest dpi) |
|---|---|---|---|
| ES, D1, stephensi, 15dpi | 36.224 | 100% (37.1) | 100% (17) |
| ES, D2, stephensi, 15dpi | Undetermined | 100% (32.7) *ave of days 7&9 | 90% (17) |
| JLW, D1, stephensi, 15dpi | 36.4231 | 100% (24.2) | 100% (17) |
| JLW, D2, stephensi, 15dpi | 36.3595 | 100% (21.0) | 100% (12), 80% (17) |
| ES, D1, gambiae 15dpi | 30.562 | 100% (46.9) | 100% (17) |
| ES, D2, gambiae 15dpi | Undetermined | 100% (18.1) | 100% (12), 92% (17) |
| ES, D1, stephensi, 16dpi | 37.038 | 100% (37.1) | 100% (17) |
| ES, D2, stephensi, 16dpi | Undetermined | 100% (32.7) *ave of days 7&9 | 90% (17) |
| JLW, D1, stephensi, 16dpi | 33.2054 | 100% (24.2) | 100% (17) |
| JLW, D2, stephensi, 16dpi | 33.3272 | 100% (21.0) | 100% (12), 80% (17) |
| ES, D1, gambiae 16dpi | 35.1362 | 100% (46.9) | 100% (17) |
| ES, D2, gambiae 16dpi | 33.8087 | 100% (18.1) | 100% (12), 92% (17) |

| | | | |
|---|---|---|---|
| NTC, water – negative control | Undetermined | NA | NA |
| Positive control – #7 2ul culture | 26.2676 | NA | NA |

**3B: Bass assay and SYBR assay qPCR comparison:**
Compared to our SYBR method described in the main text, the Bass Assay probe-based qPCR did not perform quite as well by generating more false negatives compared to SYBR and having lower sensitivity (See Table S1.5).

**Table S1.5 Bass Assay vs. SYBR assay comparison:** Cp represents the density of sporozoites in the different samples. NF54 is a strain of *P. falciparum* (*P.f.*).

| **Sample Info** | **Sample #ID** | **Bass Assay Cp** | **SYBR Assay Cp** |
|---|---|---|---|
| 100ul Plain cotton spiked with 1:10 PBS to NF54 culture, diluted | Neat | 28.0818 | 20.8941 |
| | Neat | 28.1586 | 20.6531 |
| | -1 | 31.5902 | 23.6893 |
| | -1 | 31.3893 | 23.7328 |
| | -2 | 35.4549 | 27.076 |
| | -2 | 36.1595 | 27.049 |
| | -3 | 38.2602 | 30.3946 |
| | -3 | Undetermined | **31.3999** |
| | -4 | Undetermined | Undetermined |
| | -4 | Undetermined | Undetermined |
| cotton | NTC | Undetermined | Undetermined |
| cotton | NTC | Undetermined | Undetermined |
| *An. stephensi* infected with NF54 *P.f*, 13dpbm cottons | As-8 d13 | 35.0596 | 27.1948 |
| | As-8 d13 | 36.0864 | 27.0398 |
| | As-10 d13 | 37.6146 | 30.4648 |
| | As-10 d13 | Undetermined | **30.0864** |
| | As-12 d13 | 36.1339 | 29.1585 |

| | | | |
|---|---|---|---|
| | As-12 d13 | 37.0737 | 29.009 |

| Assay | Slope | Intercept | R2 |
|---|---|---|---|
| Bass | -3.559072 | 42.388142 | 0.986662 |
| SYBR | -3.372236 | 34.041691 | 0.991813 |

While both assays detected parasites, the SYBR assay is roughly 100x more sensitive and also resulted in a better overall efficiency. Although the Bass Assay was not further optimized in our lab, it seemed that the SYBR assay performed better.

**4. Generating a standard curve and testing how sample handling processes might affect sample loss or degradation:**

*Standard curve generation*

The infected salivary glands homogenate from the dissection of 40 *An. stephensi* mosquitoes diluted in 1.2 ml 1 x PBS was estimated to contain 135,000 sporozoites per ml based on calculations scaling up from the number of sporozoites visualized in the hemocytometer counts to the volume of liquid used in the hemocytometer counting. The number of extracted sporozoites then used for the neat sample extraction for generating the standard curve is roughly 270 sporozoites per ul (in 50 ul total). This SYBR assay targets the COX1 mitochondrial gene, and each sporozoite has 20 copies of this gene (Marie et al., 2013), thus there are 5,400 copies of the gene in 1ul of the undiluted sample from this extraction. It was calculated that in 100 ul of this salivary gland homogenate that there would be 13,500 sporozoites, in 20 ul 2700 sporozoites, and in 5 ul 675 sporozoites (all multiplied by 20 for expected gene copy numbers).

Serial dilutions 1:2 a total of 8 times (1:1 through 1:128) used in the standard curve showed the expected results of a slightly better efficiency with 2 ul template samples. Overall results showed that increased template volume resulted in lower Cp values, as expected. It was determined from these data that the minimum detection threshold was lower than the 1:128 dilution, for which the Ct value was 26.11-27.62 depending on whether 1 ul or 2 ul of template was used (lower values 2 ul). Thus, it is estimated that the assay can detect <675 sporozoites. This makes sense, given that mosquitoes are thought to expectorate in the order of 10's to 100's of sporozoites at one feeding, and likely on the lower end of this range for sugar feeding, see (Churcher et al., 2017; Marie et al., 2013) and (Frischknecht et al., 2004).

*Effects of heat, time, and processing on degradation*

The effect of 24 hours at 27 °C was quite strong with most samples left at 27 °C overnight performing worse than their counterparts that were frozen immediately after their addition to either sugar soaked or dry cottons. Liquid only samples should not have had any trouble with DNA loss attributed to sample being stuck on the cotton, and indeed this was the case with most liquid samples performing the best of their set for template volume (lowest Cp). Averages are presented in Table S1.6 of samples run in duplicate. In bold are samples that weren't as close as is preferable for qPCR values since pipetting 1ul accurately can be challenging, and at low parasite numbers, the likelihood of picking up DNA from low concentrations and getting it into a reaction could be stochastic.

From these results, it can be observed that larger amounts of sporozoites added to the assay (mostly) resulted in lower Ct values as expected. There was evidence that heat contributed to sample degradation with both varieties of cotton (either treated with a sugar solution prior to use, or left dry and plain before sporozoite homogenate was added) had lower Ct scores on average when immediately frozen compared to after 24 hours in the warm and humid conditions. Liquid had lower Ct scores for every volume, suggesting that there is some loss with parasite DNA sticking to the cotton or inefficiencies in the extraction even after optimization, as would be expected. However, this loss was not severe, resulting in only a slightly lower Ct score in most comparisons.

**Table S1.6:** Samples: sporozoite standard mix from infectious *An. stephensi*, quantified 135,000 spz/ml using hemocytometer. 40 glands in 1.2 mls PBS, portioned as follows into either an Eppendorf tube (liquid samples), or onto 15 mg cotton treated either with sugar solution, or left dry "plain". Samples were either frozen immediately "frozen" or left in the 27 °C, 80% rH conditions for 24 hrs prior to extraction and qPCR SYBR assay.

| Sample volume used | Cotton with sugar | Plain cotton | Liquid (frozen) | 24hr Cotton w/ sugar | 24hr Plain cotton |
|---|---|---|---|---|---|
| 5ul | 27.1804 | 28.91635 | 25.8291 | 27.8003 | 30.3973 |
| 20ul | 26.3881 | 27.2387 | 24.35025 | **32.99555** | 29.70025 |
| 100ul | 23.2631 | 21.5206 | 21.3335 | 29.204 | 29.33915 |
