## Supplementary material for "Using a non-destructive sugar-feeding assay for sporozoite detection and estimating the extrinsic incubation period of *Plasmodium falciparum* in mosquito vectors": Supplementary data 2.pdf

**Supplementary data S2: Testing the assay at various temperatures**

**Methods**

With the goal of testing whether it was possible to use the optimized assay at various temperatures, two additional experiments were conducted.

To test lower thermal limits, 9 cups holding 6-12 *An. stephensi* mosquitoes were maintained at 27 °C for 15 days following an infectious feed then were placed in the following temperatures: (3 x 12 °C, 2 x 14 °C, 2 x 16 °C, and 2 x 18 °C). The standard optimized sugar soaked cotton assay was run from 16-21dpi with cotton collected daily from each cup, with at least one mosquito surviving in each cup during this sampling period. Samples were analyzed by Qiagen extraction and qPCR using the best SYBR methods described in the main paper. Additionally, individual *An. stephensi* mosquitoes maintained at 27 °C for 25 days post infection at which time 12 were placed at 32 °C and 12 were placed at 34 °C. All mosquitoes in both experiments were estimated to have infection prevalence of over 90% with *P. falciparum* from infectious feeds on cultured parasites. Mosquitoes in the warmer temperatures were sampled at 26 and 27 dpi. It was expected that sugar feeding at warmer temperatures would be more frequent compared to lower temperatures to maintain hydration. In lower temperatures mosquitoes would also have reduced activity levels, and, as shown from sugar feeding assays described in the main text, lower temperatures also reduce potentially sugar feeding and/or digestion rate.

**Results**

**Table S2. Comparing likelihood of sporozoite detection using this assay at various temperatures.**

| Temperature °C | Total sample number | Sporozoite positive samples (dpi) | % Positive samples |
|---|---|---|---|
| 12 °C | 18 | 0 | 0% |
| 14 °C | 12 | 2 (16, 21) | 16.67% |
| 16 °C | 12 | 1 (17) | 8.3% |
| 18 °C | 12 | 6 (17, 19, 20, 21) | 50% |
| 32 °C | 22 | 7 (26, 27) | 31.8% |
| 34 °C | 24 | 3 (26) | 12.5% |

These results show that mosquitoes kept at temperatures <16°C and >32°C were less likely to take a sugar compared to those kept between 18°C-32°C. This suggests that the use of the spit assay would be more optimal at a temperature between 18°C and 32°C and this will avoid false negative sample.
