## Supplementary material for "Using a non-destructive sugar-feeding assay for sporozoite detection and estimating the extrinsic incubation period of *Plasmodium falciparum* in mosquito vectors": Supplementary data 3.pdf

**<u>Supplementary data S3</u>: Oocyst rupture assay**

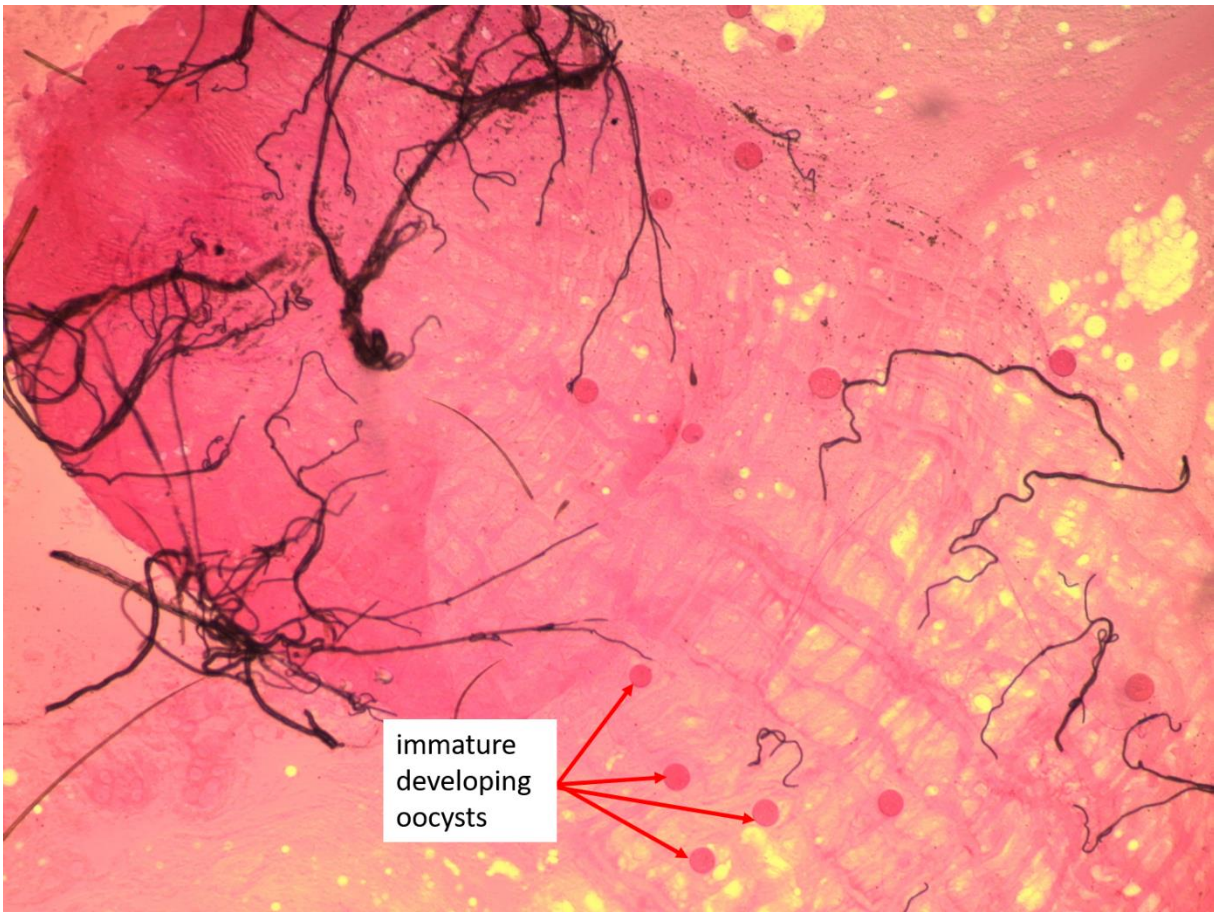


Figure S3-1: Microscopic observation (x10) of a mosquito gut dissected at 6 days post-blood-meal (dpbm) and harboring a dozen of immature developing oocysts. Mosquitoes were maintained at 27°C during parasite development. Credit: Guissou Edwige

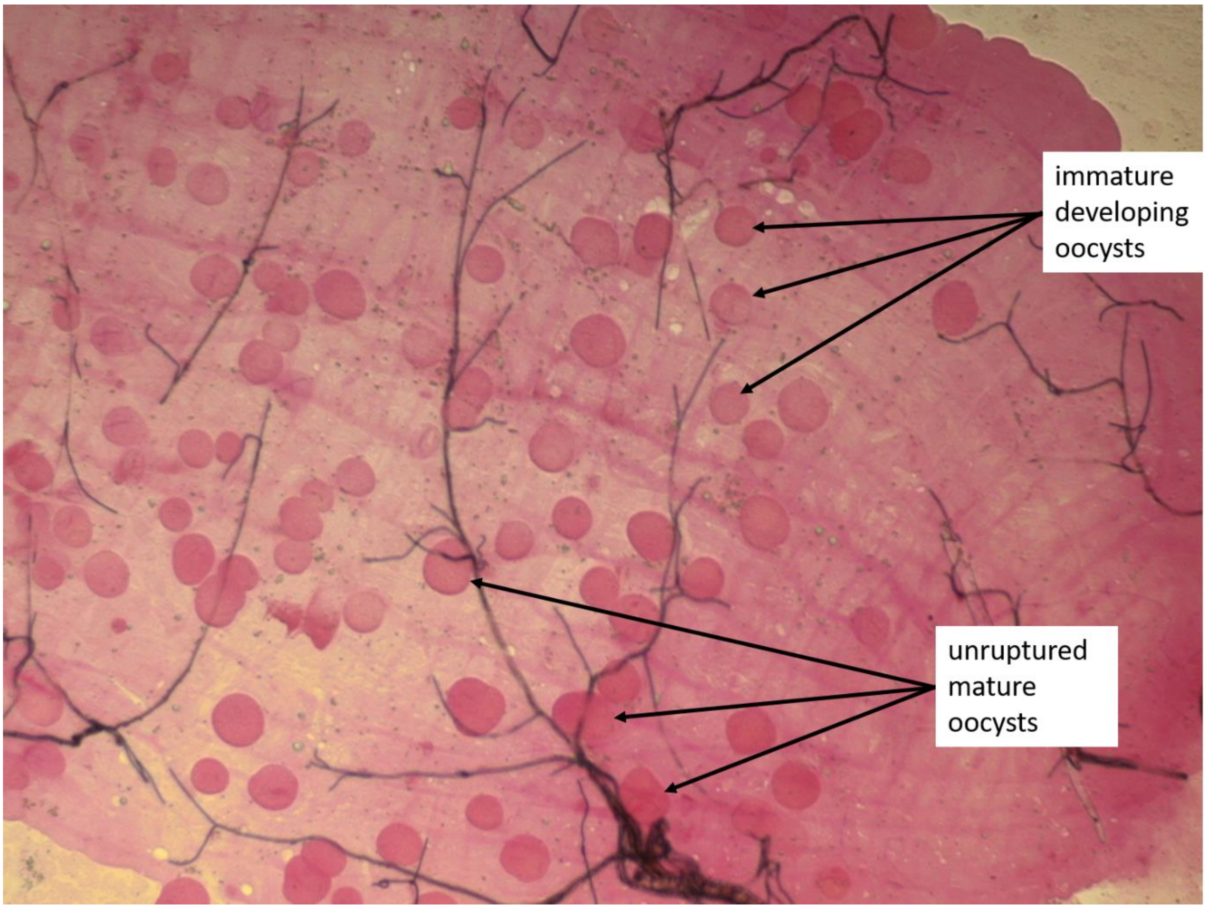


Figure S3-2: Microscopic observation (x 10) of a mosquito gut dissected at 8 days post-blood-meal (dpbm) and harboring both mature and immature oocysts. Note the protrusion of the capsule of some oocysts that may here result from the application of the coverslip. In our experiment, these distorted oocysts were recorded as intact unruptured oocysts because the capsule has not broken yet. Mosquitoes were maintained at 27°C during parasite development. Credit: Guissou Edwige

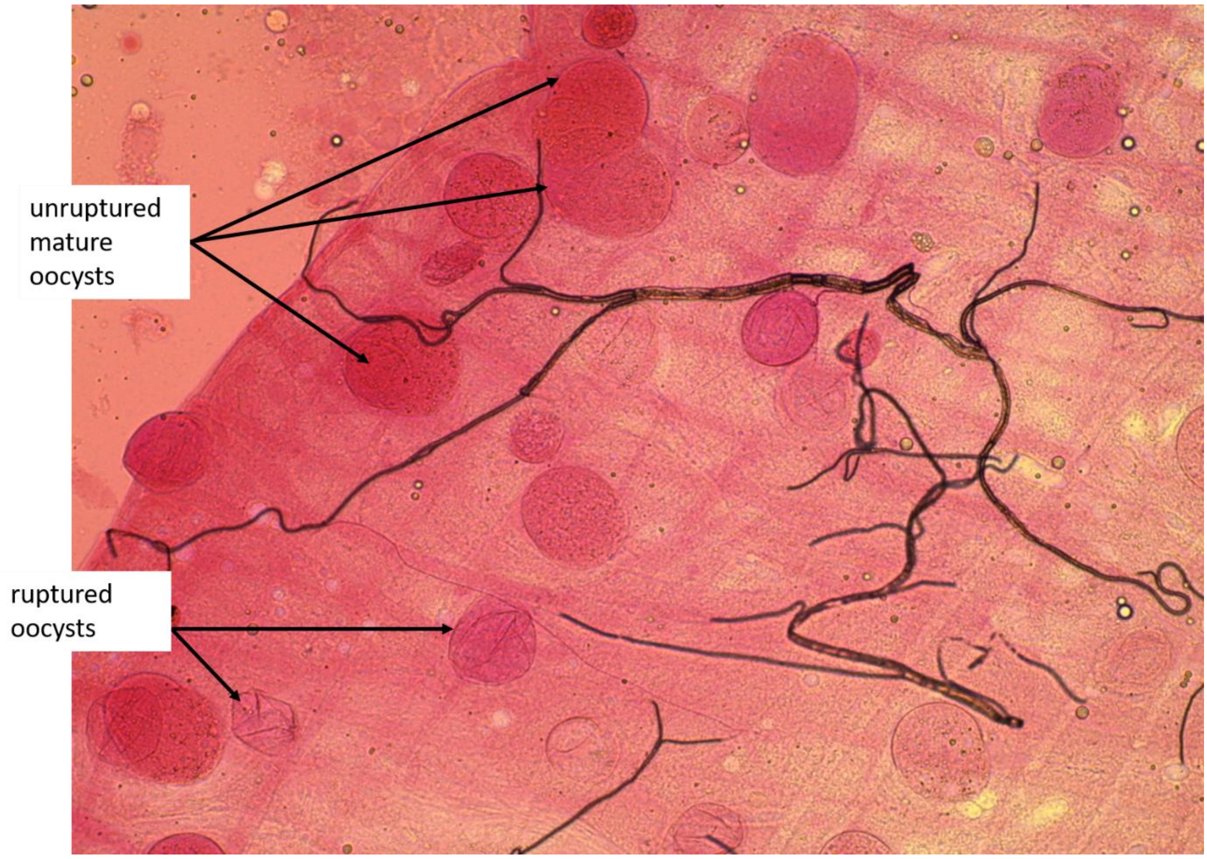


Figure S3-3: Microscopic observation (x 20) of a mosquito gut dissected at 12 days post-blood-meal (dpbm) and harboring both ruptured and unruptured mature oocysts. The ruptured oocysts are characterized by empty and withered capsules. Ruptured oocysts look like a hatched egg from which only the shell remains. Mosquitoes were maintained at 27°C during parasite development. Credit: Guissou Edwige

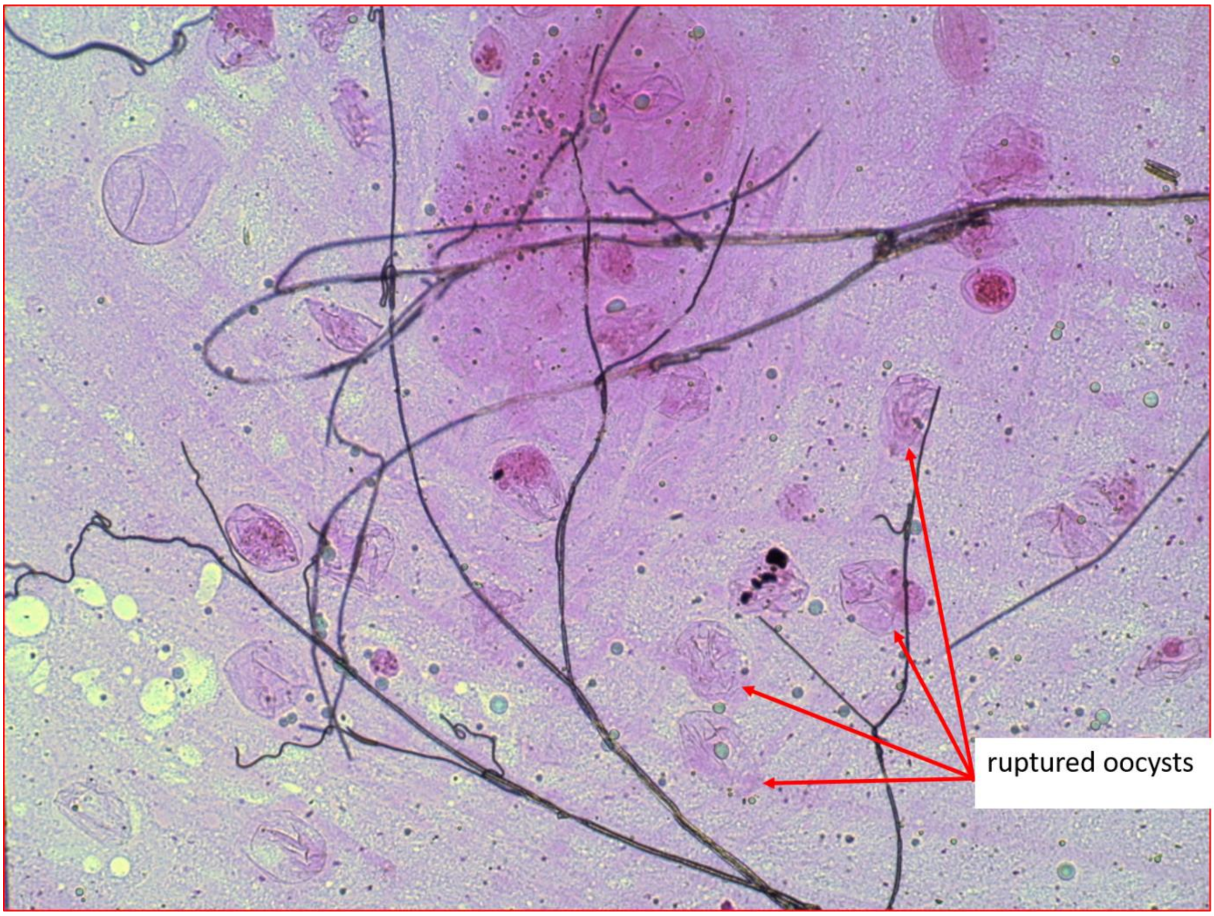


Figure S3-4: Microscopic observation (x 20) of a mosquito gut dissected at 12 days post-blood-meal (dpbm) and harboring ruptured oocysts only. The ruptured oocysts are characterized by empty and withered capsules. Ruptured oocysts look like a hatched egg from which only the shell remains. Mosquitoes were maintained at 27°C during parasite development. Credit: Guissou Edwige
