## Supplementary material for "Using a non-destructive sugar-feeding assay for sporozoite detection and estimating the extrinsic incubation period of *Plasmodium falciparum* in mosquito vectors": Supplementary data 4.pdf

**<u>Supplementary data S4</u>: Infection level and mosquito survivorship in experiment 1**

The proportion of infected mosquitoes (prevalence) in isolate A (95% (20/22)) and in isolate B (100% (20/20)) was statistically similar (GLM binomial: $LRT\ X^2_1 = 1.31$, P = 0.25, Figure S4A). The mean number (± se) of developing oocysts in infected females (density) was significantly higher for isolate B (191.65 ± 21) than for A (13.86 ± 2) (GLM negative binomial: $LRT\ X^2_1 = 133$, P < 0.001, Figure S4B). Although the prevalence of sporozoites based on microscopic observation from 14 to 16 dpbm was similar for both parasite isolates (A: 95% (37/39); B: 97% (36/37); $LRT\ X^2_1 = 0.30$, P = 0.58, Figure S4C left panel), that based on qPCR was slightly higher for isolate B (100% (37/37)), than for A (92% (36/39) ($LRT\ X^2_1 = 4.12$, P = 0.04, Figure S4C). The sporozoite density based on the scores was higher for isolate B (median=3) than for A (median=2), ($LRT\ X^2_1 = 1.65$, P < 0.001, Figure S4D), thus confirming the oocyst observation. However, the sporozoite density based on qPCR (Ct) was similar for both parasite isolates ($LRT\ X^2_1 = 0.004$, P = 0.53, Figure S4D). Mosquito survival was not significantly associated to parasite isolates (survival cox model: (LRT $X^2_1$=0.54, P=0.46. figure S4E).

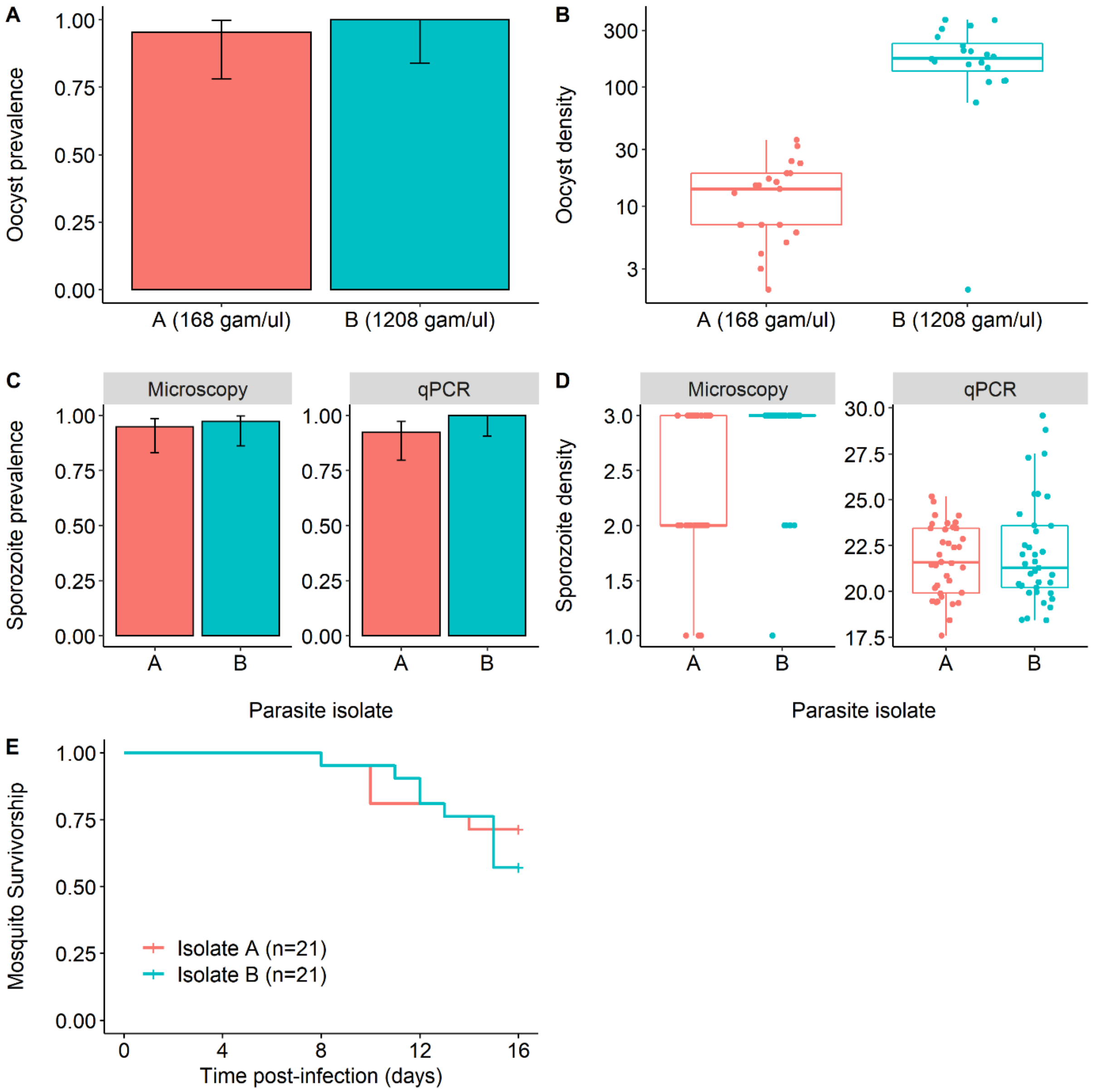


**Figure S4: Infection level at two distinct time points over the course of infection (oocyst and sporozoite stages).** A: Oocyst prevalence (± 95% CI) on day 8-9 post-blood meal (dpbm), expressed as the number of mosquito females harboring at least one oocyst in their midguts out of the total number of dissected females, for each parasite isolate (red bar: isolate A, blue bar: isolate B). B: Oocyst density at 8-9 dpbm, expressed as the mean number of developing oocysts (± se) in the guts of infected females, for each parasite isolate. The different numbers in parentheses in Figures 1A and 1B indicate gametocymia (number of gametocytes / ul of blood) for isolates A and B. C: Sporozoite prevalence (± 95% CI) at 14-16 dpbm, expressed as the number of mosquito salivary glands detected positive to *P. falciparum* using microscopic observation (left panel) or qPCR (right panel) out of the total number of dissected salivary glands, for each parasite isolates. D: Sporozoite density at 14-16 dpbm, expressed as the median score assigned to each salivary gland (1: a few sporozoites, 2: a moderate level of sporozoites, 3: numerous sporozoites) when using microscopy for parasite detection or the median Ct values

when using qPCR (the lower the Ct, the higher the sporozoite density) for each parasite isolate.
E: Survival of mosquito females used to collect saliva for each parasite isolates.
