## Supplementary material for "Using a non-destructive sugar-feeding assay for sporozoite detection and estimating the extrinsic incubation period of *Plasmodium falciparum* in mosquito vectors": Supplementary data 5.pdf

**Supplementary data S5:** Infection level and mosquito survival in experiment 2

The parasite load (inferred from the Cp values: the higher the Cp, the lower the parasite load) in infected *An. coluzzii* carcasses was lower than that of *An. gambiae* and *An. arabiensis* (mean Cp ± se: 26.64 ± 1.24, 23.6 ± 0.86 and 21.51 ± 0.6, respectively, KW $X^2_2 = 9.3$, P=0.009, figure S5.1A). In addition, there was a positive relationship between the probability to generate *P. falciparum* positive cotton samples and infection intensity in individual females (LRT $X^2_1$=10, P=0.002, figure S5.1B).

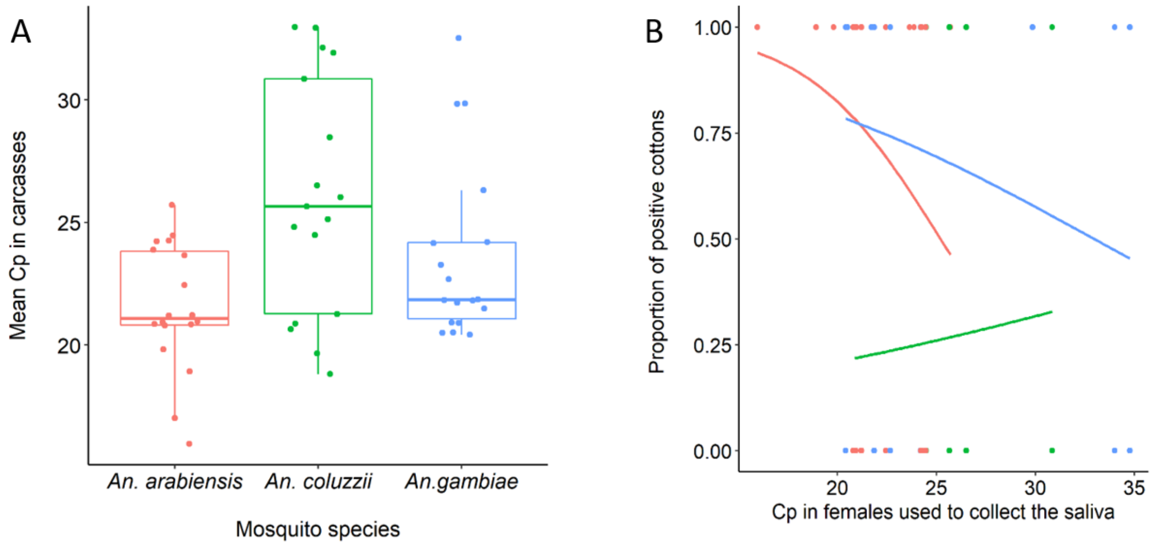


**Figure S5.1.** Infection level. (A) Infection load in mosquito females used to collect saliva (expressed as the mean Cp values from the qPCR output. The lower the Cp, the higher the parasite load) for each anopheline species (red: *An. arabiensis*, *green: An. coluzzii*, blue: *An. gambiae)*. (B) Estimated probability of *P. falciparum* carriage in cottons as a function of infection load in females used depositing saliva on these cottons. The lower the Cp, the higher the infection load. The red, blue and green lines show the relationship for *An. arabiensis*, *An. coluzzii* and *An. gambiae* respectively.

*An. arabiensis* survived better in plastic tubes than both *An. coluzzii* and *An. gambiae* (survival cox analysis on infected and uninfected individuals (n=60): LRT $X^2_2$=7.8. P=0.02, median survivorship: 18.5, 12 and 12.5 days, in *An. arabiensis*, *An. coluzzii* and *An gambiae*, respectively, figure S5.2).

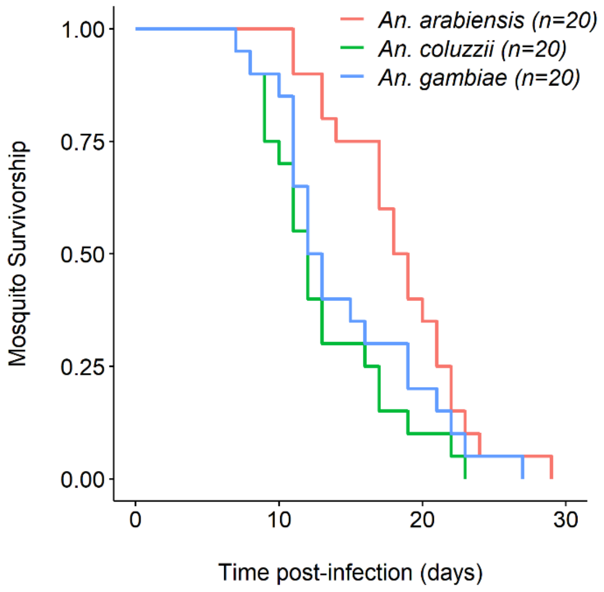


**Figure S5.2.** Survival of mosquito females used to collect saliva for each anopheline species.
