## Supplementary material for "Using a non-destructive sugar-feeding assay for sporozoite detection and estimating the extrinsic incubation period of *Plasmodium falciparum* in mosquito vectors": Supplementary data 6.pdf

**Supplementary data S6**: Relationship between mosquito sugar feeding and *P. falciparum* positivity in cottons

Table 1: Evaluation of the presence colored fecal dots from 14 to 24 days after the infectious blood meal (dpbm) from 16 females infected with *P. falciparum*. Each id represents a female. Color_dpbm represents the day of cotton collection and the observation (1) or not (0) of colored fecal dots. The death column corresponds to the day the mosquito died. Red color: *P. falciparum* positive sample, Blue color: *P. falciparum* negative sample, NA: not available sample

| Id | color_dpbm14 | color_dpbm15 | color_dpbm16 | color_dpbm17 | color_dpbm18 | color_dpbm19 | color_dpbm20 | color_dpbm21 | color_dpbm22 | color_dpbm23 | color_dpbm24 | Death(dpbm) |
|---|---|---|---|---|---|---|---|---|---|---|---|---|
| 2 | 0 | 0 | 1 | 1 | NA | NA | NA | NA | NA | NA | NA | 18 |
| 4 | 0 | 0 | 1 | 0 | 1 | 0 | 0 | NA | NA | NA | NA | 21 |
| 6 | 1 | 0 | 0 | NA | NA | NA | NA | NA | NA | NA | NA | 16 |
| 7 | 1 | 1 | 1 | NA | NA | NA | NA | NA | NA | NA | NA | 17 |
| 10 | 1 | 1 | 1 | 1 | 1 | NA | NA | NA | NA | NA | NA | 19 |
| 11 | 1 | 1 | 1 | 1 | NA | NA | NA | NA | NA | NA | NA | 18 |
| 13 | 1 | 1 | 1 | 1 | 1 | 1 | 1 | 1 | 1 | 1 | 1 | 24 |
| 15 | 0 | 1 | 1 | 1 | 1 | 0 | 1 | 1 | 1 | 1 | NA | 23 |
| 16 | NA | 0 | 0 | 1 | 1 | NA | NA | NA | NA | NA | NA | 18 |
| 18 | 0 | 0 | 1 | 1 | 1 | 1 | 0 | NA | NA | NA | NA | 21 |
| 19 | 1 | 1 | 1 | 1 | 1 | 1 | 0 | NA | NA | NA | NA | 21 |
| 20 | 1 | 1 | 1 | 1 | 0 | 0 | NA | NA | NA | NA | NA | 20 |
| 21 | 0 | 0 | 0 | 0 | 1 | 0 | 0 | 0 | 0 | 0 | 1 | 24 |
| 23 | 0 | 1 | 1 | NA | NA | NA | NA | NA | NA | NA | NA | 17 |
| 26 | 1 | 1 | NA | NA | NA | NA | NA | NA | NA | NA | NA | 15 |
| 30 | 1 | 1 | 1 | NA | NA | NA | NA | NA | NA | NA | NA | 16 |

This table shows that the probability of detecting *P. falciparum* positive cottons was not related to the detection of colored fecal dots on the papers. We must therefore continue the investigations in order to know the frequency of sugar feeding intake of mosquitoes because this will improve the efficiency of the spit assay.
